# Osteoclast-Cancer Cell Metabolic Symbiosis Renders PARP Inhibitor Therapy Resistance in Bone Metastasis

**DOI:** 10.1101/2023.08.15.553338

**Authors:** Huijuan Fan, Zhanao Xu, Ke Yao, Bingxin Zheng, Yuan Zhang, Xuxiang Wang, Tengjiang Zhang, Xuan Li, Haitian Hu, Bin Yue, Zeping Hu, Hanqiu Zheng

**Affiliations:** Department of Basic Medical Sciences, School of Medicine, Tsinghua University, Beijing, 100084, China; School of Pharmaceutical Sciences, Tsinghua University, Beijing, 100084, China; Department of Orthopedic Oncology, the Affiliated Hospital of Qingdao University, No. 59 Haier Road, Qingdao, Shandong, 266000, China; State Key Laboratory of Molecular Oncology and Department of Basic Medical Sciences, School of Medicine, Tsinghua University, Beijing, 100084, China

## Abstract

Seventy percent of patients with late-stage breast cancer develop distal bone metastases; however, the mechanism by which the metabolic microenvironment affects resistance to therapy remains unknown. We investigated the metabolic bone microenvironment and identified glutathione metabolism as the top pathway in osteoclasts, which provides feedback to tumor cells to help neutralize oxidative stress and generate PARP inhibitor (PARPi) therapy resistance. GPX4, the critical enzyme responsible for glutathione oxidation, was upregulated during PARPi therapy through stress-induced ATF4-dependent transcriptional programming. The increased absorption of glutamine and the upregulation of GPX4 expression work in concert to enhance glutathione metabolism in cancer cells. Human clinical sample analysis of paired primary breast tumor and bone metastasis samples revealed that GPX4 was significantly induced in bone metastases. Combination therapy utilizing PARPi and zoledronate, which blocks osteoclast activity and thereby reduces the microenvironmental glutamine supply, generates a synergistic effect in reducing bone metastasis. Thus, our results identified an essential metabolic symbiosis between bone-resident cells and metastatic cancer cells during PARPi therapy.

**SIGNIFICANCE:** Osteoclast-derived glutamine is taken up by tumor cells to synthesize glutathione and neutralize the ROS generated by PARPi. This is the first example of “metabolic symbiosis” in therapeutic resistance of bone metastasis.

## INTRODUCTION

A large proportion of patients with late-stage breast cancer develop bone metastasis. These patients suffer from intolerable bone pain, bone fracture, and sometimes life-threatening hypercalcemia(1). Bone metastasis might also be a reservoir of “tumor seeds”, which colonize other distal organs to generate systematic metastasis(2). However, many current therapies against bone metastasis are palliating(1), without significant extension of patient survival time, as bone metastases often resist these therapies.

The progression of bone metastasis is heavily influenced by the tumor microenvironment(3–6), with stromal cells such as osteoclasts playing a major role in late-stage osteolytic bone metastasis(6). Through several well-defined signaling molecules, including parathyroid hormone-related protein (PTHrP), receptor activator of nuclear factor kappa-Β ligand (RANKL), Jagged1 and others, tumor cells promote the formation of multi-nucleated, bone resorptive osteoclasts(7–9). The resulting degradation of bone releases growth factors and cytokines from the bone matrix, which in turn provide feedback to tumor cells and promoting metastatic outgrowth. This is known as the “vicious cycle” of osteolytic bone metastasis. Some therapeutic agents, such as zoledronate and RANKL-neutralizing antibody (denosumab), have been approved to treat bone metastasis by blocking this cycle. In addition to these anti-osteolytic therapies, conventional DNA-damaging chemotherapies have also been used. Recently, poly (ADP-ribose) polymerase inhibitor (PARPi) was approved for breast and ovarian cancer based on its synthetic lethality in inducing DNA double-strand breaks (DSBs) and cell death in BRCA1/2-mutant patients or in other DNA repair deficient patients(10–14). However, the therapeutic effect of PARPi in bone metastasis and possible resistance mechanisms are not yet well understood(15).

Bone stromal cells are critical to the development of therapy resistance. For example, during chemotherapy, the adjacent stromal cell niche is damaged and rebuilt by engrafted leukemic cells, the regenerated bone niche protects tumor cells from death(16). Our previous studies revealed that osteoblastic Jagged1 helps bone metastatic tumor cells survive chemotherapy(5). In addition, the expansion of pericytes in bone generates a proliferative microenvironment that is resistant to radiation and chemotherapy(17). In recent years, aberrant metabolic reprogramming has become a hallmark of cancer progression and metastasis(18–20). For example, microenvironment-derived pyruvate helps shape the lung niche to promote breast cancer lung metastasis(21). One common hurdle that tumor cells must overcome during metastasis progression and DNA damaging-based therapy is that cancer cells urgently need to cope with increased reactive oxidative species (ROS) to survive. Reports suggest that disseminated tumor cells face increased ROS at distal organ sites, and only when tumor cells cope with this increased ROS level by shunting their metabolic pathways can they survive and generate metastatic colonies(22). DNA-damaging agents are known to cause cell death partially by inducing ROS production(24). Thus, it is plausible that cancer cells that successfully adopt a metastatic niche might already become extremely resistant to chemotherapy due to this pre-evolved metabolic adaptation. However, metabolic reprogramming of bone stromal cells and metabolic crosstalk between stromal and tumor cells during bone metastasis and therapy resistance have yet to be fully investigated.

Osteoclasts, which are large, multinucleated cells generated from the fusion of their progenitor cells (macrophages), produce significant amounts of acid and proteases to help degrade the bone. However, it is unknown whether hyperactivation and an increased number of osteoclasts generate metabolic symbiosis between osteoclasts and tumor cells, especially under stress conditions induced by cancer therapy. Our current study used metabolic profiling and functional analysis to investigate this possibility and found that mature osteoclasts provide glutamine to tumor cells. In turn, tumor cells upregulate essential transporter and enzyme activity to enhance the absorption of these amino acids and upregulate glutathione metabolism. Glutathione is essential for tumor cell survival during PARPi therapy, as it is utilized as a major reducing agent to neutralize reactive oxidative species (ROS). Our findings suggest that this metabolic symbiosis between tumor cells and osteoclasts could be the Achilles heel in treating patients with bone metastatic disease.

## RESULTS

### Metabolic Profiling in Osteoclast Maturation

To understand the potential role of osteoclasts in therapy resistance during bone metastasis, we followed the experimental design illustrated in Fig. 1A. Briefly, bone marrow cells from the hindlimbs of 6-8 weeks old mice were collected and cultured *in vitro* in the presence of macrophage colony-stimulating factor (M-CSF) and RANKL to induce osteoclastogenesis. Conditioned media (CM) from pre-osteoclasts (macrophages) and mature osteoclasts were collected. These CM were filtered through a 3K-centrifugal filter column (molecules with molecular weight larger than 3,000 Daltons will not flow through the filter) to remove majority of protein contents. The CM were then used in cell-killing assays with different chemotherapeutic agents. Osteoclast maturation was confirmed using tartrate-resistant acid phosphatase (TRAP) staining and RNA-sequencing (RNA-seq) to examine the expression of essential osteoclast marker genes (Fig. 1B and C and Supplementary Fig. S1A). Bone metastatic tumor cells were cultured in CM and treated with the respective therapeutic agents to determine the potential therapeutic resistance caused by osteoclast CM. In both 4T1.2 and SCP28 cells, a mouse and a human bone metastasis-prone mammary tumor cell line(26,27), the addition of osteoclast CM generated a strong survival effect when cancer cells were treated with olaparib or cisplatin (Fig. 1D). In contrast, when cells were treated with paclitaxel or vincristine, there was no survival benefit from osteoclast CM (Supplementary Fig. S1B). Cisplatin acts as a DNA-damaging agent by crosslinking the purine bases on the DNA to form DNA adducts(28), while paclitaxel and vincristine act as cytotoxic agents by stabilizing or destabilizing microtubules(29–31). Thus, our results suggest that osteoclast CM generates therapeutic resistance specifically in DNA damaging-based therapies, but not in other types of therapies. Since the CM had been filtered to remove any molecule larger than 3,000 Daltons, it is unlikely that the resistant phenotype was caused by proteins in the OC CM.

**Figure. 1.**
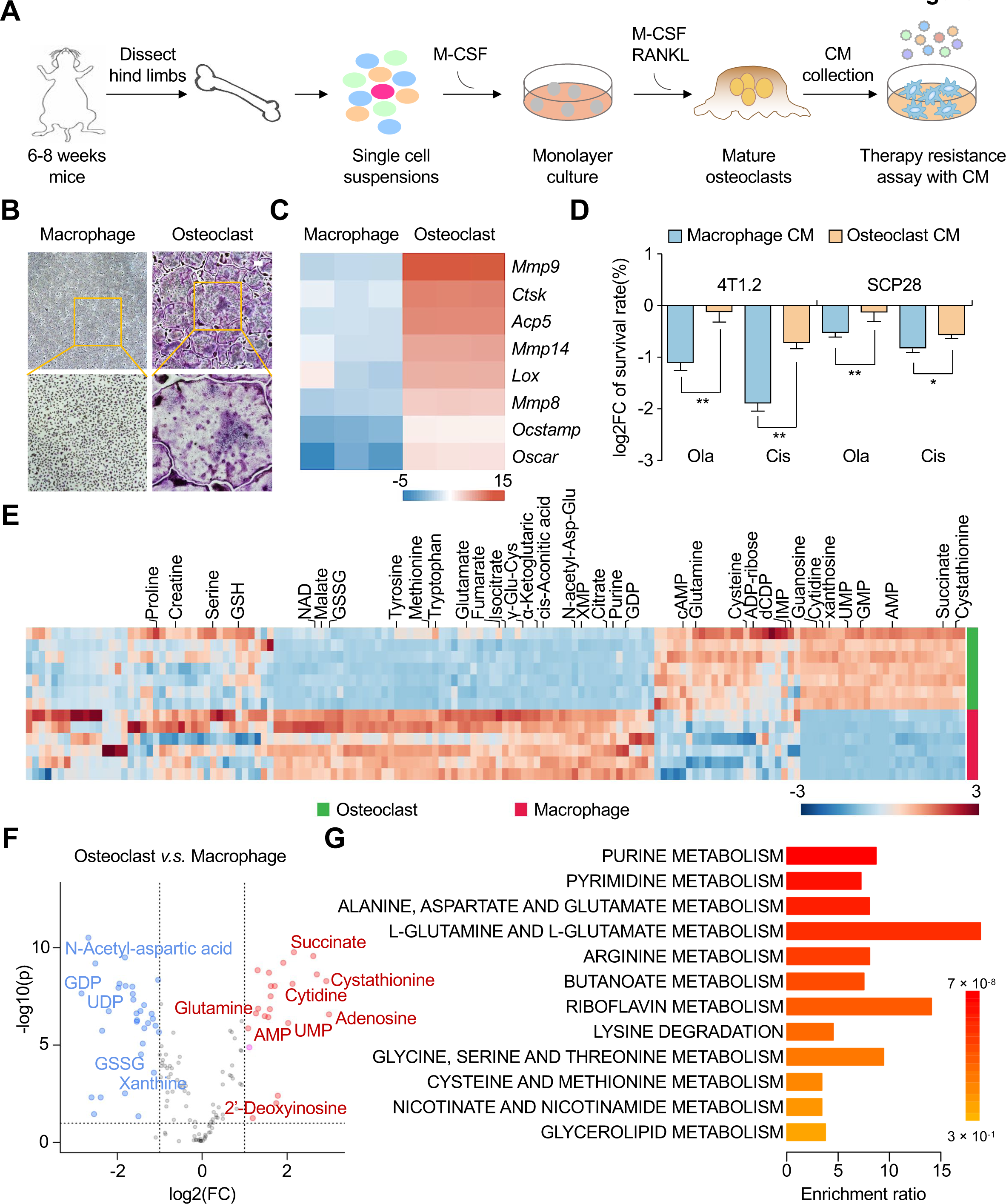
Metabolic profiling of mature osteoclast and its precursor macrophage. (**A**) The workflow of *in vitro* osteoclastogenesis assay for conditioned media collection and the testing of therapeutic resistance. (**B**) Osteoclasts were differentiated based on experimental procedure in A. The cell culture was then stained for TRAP-positive cells. The representative images of TRAP staining were shown. Scale bar, 200 μm. (**C**) RNA-seq was performed by using RNAs from macrophages and mature osteoclasts. The differentially expressed osteoclast marker genes were presented in a heatmap. (**D**) The survival rates of 4T1.2 cells (left) and SCP28 cells (right) treated with either DMSO, olaparib (25 μM), or cisplatin (3 μM) in the presence of macrophage CM or osteoclast CM mixed with regular culture medium at the ratio of 1:1. (**E**) Heatmap of metabolites in six replicates of macrophage samples and seven osteoclast samples as determined by mass-spectrometry. Macrophage: n = 6; Osteoclast: n = 7. (**F**) Volcano plots of differentially presented metabolites between macrophages and osteoclasts. Each dot represents one metabolite. Metabolites used in this analysis were changed at least two folds and with false discovery rate (FDR) less than 0.05. (**G**) Metabolic pathway enrichment analysis of increased metabolites in osteoclasts *v.s.* macrophages. Metabolites used in this analysis were changed at least two folds and with false discovery rate (FDR) less than 0.05. **See also Supplementary Figure. S1.**

Recent studies have begun to appreciate the role of metabolites in cancer therapy resistance(18,19,32,33). There might be some metabolites in CM from OC which are responsible for the observed therapy resistance. To systematically profile the metabolomes of mature osteoclasts *v.s.* their precursors, macrophages and osteoclasts were lysed for targeted metabolomics analysis(34). A principal component analysis was performed based on the detected metabolites. All replicates of the metabolome from osteoclasts were well separated from their macrophage precursors, confirming the robustness of the metabolome profiling (Supplementary Fig. S1C). Osteoclasts demonstrated clear metabolic reprogramming compared with macrophages, as many essential amino acids and nucleotides were altered during osteoclastogenesis (Fig. 1E and F and Supplementary Fig. S1D). Metabolic pathway analysis revealed that nucleotide biosynthesis, pentose phosphate pathway, tricarboxylic acid cycle (TCA cycle), and amino acid metabolism, especially alanine, aspartate, and glutamine metabolism, were enriched in osteoclasts (Fig. 1G and Supplementary Fig. S1E). A few examples of these critical pathways, including glutamine, glutathione (GSH), isocitrate, and succinate, were significantly upregulated in mature osteoclasts. Simultaneously, glutathione disulfide (GSSG), citrate, cis-aconitic acid, and α-ketoglutarate (α-KG) levels were significantly decreased in osteoclasts (Supplementary Fig. S1F).

### Glutamine Pathway is Essential for Tumor Cell Survival during Treatments of DNA-damaging Agents

To examine whether specific metabolic pathways or enzymes are involved in therapy-resistance to DNA-damaging agents, we utilized a PARP inhibitor as a model insult to the DNA stability of tumor cells. MDA-MB-231, a human breast cancer cell line, was treated with either the DMSO control or olaparib (a type of PARPi), and mRNA was collected for RNA-seq. The upregulated genes in the PARPi-treated group *v.s.* the control group were matched with a metabolic gene set based on the Molecular Signature Database(35–37) and Kyoto Encyclopedia of Genes and Genomes (KEGG) to identify metabolism-related genes that were significantly upregulated.

Gene set enrichment analysis (GSEA) was performed to detect enriched metabolic pathways. The most significantly enriched pathway was cellular response to stress (Fig. 2A). The top 20 upregulated metabolic-related genes included indoleamine 2,3-dioxygenase 1 (*IDO1*) and serine dehydratase (*SDS*), which are enzymes related to tryptophan and serine metabolism, respectively (Fig. 2B). We compared the top upregulated metabolites in osteoclast differentiation, the associated metabolites in the cellular response to stress, and metabolites associated with highly changed genes from RNA-seq to generate a list of candidate metabolites that might play a role in PARPi therapy resistance (Fig. 2C and D). To functionally test the role of these metabolites in PARPi therapy resistance, we supplemented the culture media with different concentrations of individual metabolites during PARPi treatment. In both 4T1.2 and SCP28 cells, the addition of succinic acid, L-serine, L-tryptophan, cytidine, and cystathionine had little effect on tumor cell survival in PARPi therapy (Supplementary Fig. S2A-J). In contrast, higher concentrations of glutamine enhanced cell survival in both the cell lines (Fig. 2E and F).

**Figure. 2.**
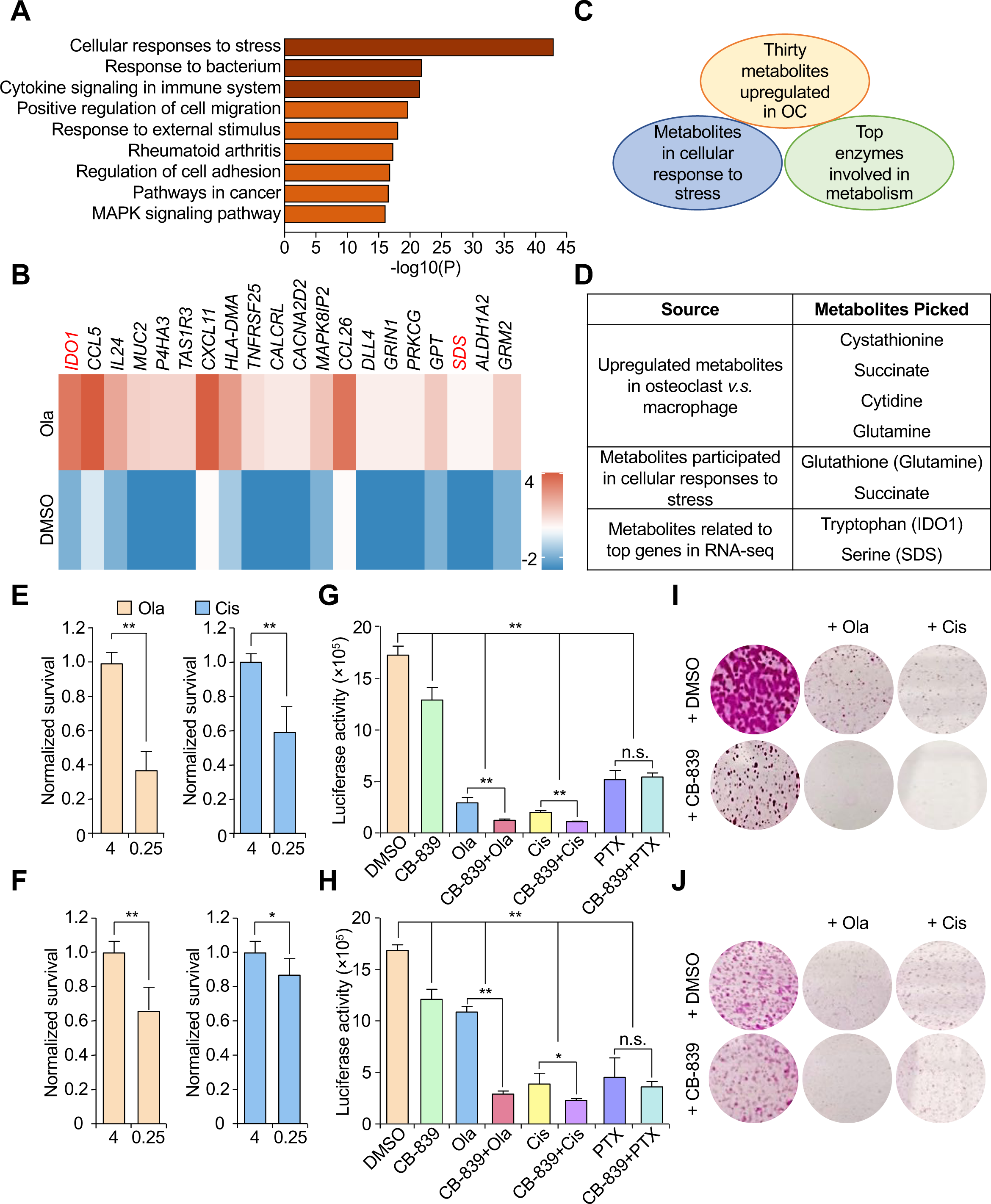
Glutamine metabolism promotes cancer cell PARPi therapy resistance. (**A**) MDA-MB-231 cells were treated with DMSO or olaparib (40 µM) for 3 days. Cells were collected and RNA was extracted for RNA-seq analysis. The upregulated genes in the olaparib treatment group were matched with a metabolic gene set based on the Molecular Signature Database and the KEGG of Genes and Genomes database. Gene set enrichment analysis (GSEA) was performed to detect enriched metabolic pathways. (**B**) Heatmap of top 20 metabolic genes upregulated in the MDA-MB-231 cell line under olaparib *v.s.* DMSO treatment from experiment performed in A. (**C**) Schematic representation of the method of selecting potential PARPi therapy resistance metabolites. Thirty metabolites were upregulated in osteoclasts *v.s.* macrophages, metabolites upregulated in the cellular response to stress, and the top 20 enzymes involved in metabolism were potential candidate metabolites. Additional criteria, such as tumor cell permeability and limited supply in the tumor microenvironment, were also considered. (**D**) The candidate PARPi resistance metabolites were selected for further functional analysis. (**E-F**) Normalized survival rates of 4T1.2 and SCP28 cells treated with olaparib (50 μM) or cisplatin (10 μM). DMSO was used as a control. Cells were cultured in either 4 mM or 0.25 mM extracellular glutamine. (E) 4T1.2 cells; (F) SCP28 cells. (**G-H**) 4T1.2 and SCP28 cells were treated with the indicated concentrations of CB-839, olaparib, cisplatin, paclitaxel, or the indicated combination treatment. CB-839 is a small molecule inhibitor of GLS1. (G) 4T1.2 cells; (H) SCP28 cells. (**I-J**) Colony formation assay of 4T1.2 and SCP28 cells treated with the indicated concentrations of olaparib, cisplatin, or combination therapy with CB-839. (I) 4T1.2 cells; (J) SCP28 cells. Ola, olaparib; Cis, cisplatin; PTX, paclitaxel. Data are presented as mean ± SEM. “*n.s.”* means not significant. *p < 0.05, **p < 0.01 by Student’s t-test (A, F, G, and H). **See also Supplementary Figure. S2.**

Extracellular glutamine is transported into the cells through a specific cell surface transporter, Solute Carrier Family 1 member 5 (SLC1A5). Once in the cell, glutamine is catalyzed by glutaminase 1/2 (GLS1/2) to generate glutamate for further utilization (Supplementary Fig. S2K). GSH, a downstream metabolite of glutamine, is one of the major molecules in balancing the redox pathway to support tumor cell survival during chemotherapies(38). Consistent with this idea, treatment of small molecule inhibitor (CB-839) against GLS1 synergized with PARPi treatment to kill both 4T1.2 and SCP28 cells (Fig. 2G-J). We then tested the potential role of the glutamine pathway in two additional commonly used chemotherapeutic agents for patients with breast cancer: cisplatin and paclitaxel. Again, cisplatin, but not paclitaxel, worked synergistically with CB-839 to kill both the cell lines (Fig. 2G-J). These results suggest that an intact glutamine metabolism pathway may promote cell survival only during DNA-damaging therapy.

To complement the above pharmacological inhibitor studies, we then genetically knocked down (KD) GLS1 to block the use of glutamine in SCP28 and 4T1.2 cells by two independent shRNA constructs and performed functional analysis (Supplementary Fig. S3A). The KD of GLS1 sensitized tumor cells to olaparib or cisplatin treatment (Fig. 3A and B). KD of GLS1 also increased γH2AX expression, an indication of increased DNA double-strand break (DSB) damage within tumor cells (Fig. 3C and D). To examine whether glutamine metabolism is critical to bone metastasis *in vivo*, we intracardially (IC) injected either vector control or GLS1-KD 4T1.2 cells into BALB/c mice. Bone metastatic burden was monitored using *in vivo* bioluminescence imaging (BLI). GLS1 KD decreased bone metastatic progression (Supplementary Fig. S3B and C).

**Figure. 3.**
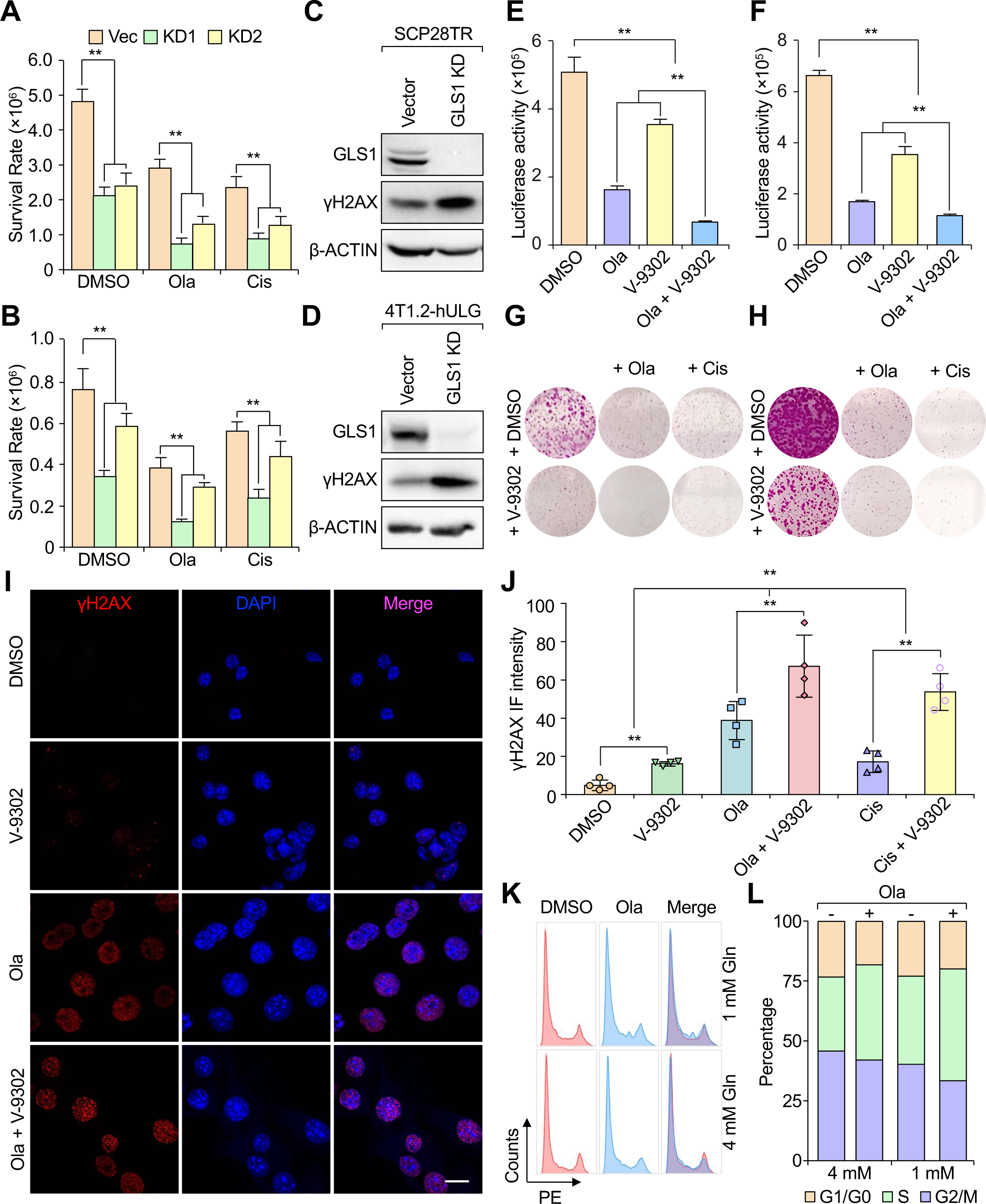
Extracellular glutamine promotes cancer cell survival during PARPi therapy. (**A-B**) Survival rates of vector control and GLS1 KD cells (two shRNAs) treated with either DMSO, olaparib (25 μM), or cisplatin (3 μM). (A) SCP28 cells. (B) 4T1.2 cells. (**C**) The expression level of γH2AX protein in SCP28-vector or in SCP28-GLS1 KD cells was determined by immunoblotting. β-ACTIN was used as internal loading control. (**D**) The expression level of γH2AX protein in 4T1.2-vector control or in 4T1.2-GLS1 KD cells was determined by immunoblotting. β-ACTIN was used as internal loading control. (**E-F**) SCP28 cells or 4T1.2 cells were treated with either DMSO control, olaparib, V-9302 or the indicated combination therapy. SCP28 cells in E. 4T1.2 cells in F. (**G-H**) Colony formation assay of SCP28 or 4T1.2 cells treated with DMSO control, olaparib, cisplatin, V-9302 or the indicated combination therapy. SCP28 cells in G and 4T1.2 cells in H. (**I**) 4T1.2 cells were treated with DMSO control, olaparib, V-9302 or the indicated combination therapy. 2 days later, cells were fixed and IF stained with γH2AX (red). Nuclei were counter-stained with DAPI (blue). Scale bar: 20 μm. (**J**) Quantification of the staining intensity of γH2AX from experiment in I and in Supplementary Fig. S3D. (**K**) Flow cytometry analysis of cell cycle of SCP28 cells treated with either DMSO or olaparib in normal (4 mM) or low (1 mM) glutamine media. (**L**) Quantification of cell cycle phases in K. “-” indicates DMSO-treated group, “+” indicates olaparib-treatment group. Data represents mean ± SEM. **p < 0.01 by Student’s t-test (A, B, E, F, and J). **See also Supplementary Figure. S3.**

### Extracellular Glutamine Promotes Cancer Cell Survival during PARPi Therapy

It is worth to know that tumor cells mostly rely on an extracellular rather than self-generated supply of glutamine to survive and proliferate(39–42). Interestingly, although glutamine is abundant in blood circulation, its quantity is extremely limited in the tumor microenvironment(43–46). We reasoned that bone metastatic tumor cells might rely on osteoclast-derived glutamine to survive the olaparib therapy. To understand the importance of extracellular glutamine in therapy resistance, we blocked the glutamine transporter SLC1A5 using a small-molecule compound (V-9302) and examined its effect on cell survival during PARPi treatment. When V-9302 and olaparib were simultaneously applied to tumor cells, they synergistically caused tumor cell death in both SCP28 and 4T1.2 cells (Fig. 3E-H). Similarly, V-9302 and cisplatin synergistically induced tumor cell death (Fig. 3G and H). Since the mechanism of action of PARPi is to generate DNA DSBs, we examined the level of DSBs using immunofluorescence (IF) staining of γH2AX. As expected, treatment with PARPi increased the expression of γH2AX in 4T1.2 cells. In contrast, treatment with V-9302 modestly increased γH2AX levels. Interestingly, combined treatment with both PARPi and V-9302 led to a striking upregulation of γH2AX (Fig. 3I and J). Similarly, combination therapy with V-9302 and cisplatin synergistically increased the expression level of γH2AX in tumor cells (Fig. 3J and Supplementary Fig. S3D), whereas combination therapy with paclitaxel and V-9302 had little effect on γH2AX levels (Supplementary Fig. S3E). To examine cellular apoptosis status, IF staining of cleaved caspase-3 (CC3) was performed. There was almost no detectable CC3 staining in the DMSO group and only modest CC3 staining in olaparib or V-9302 singly treated cells, whereas a substantial increase in CC3 staining was observed in the combination therapy group (Supplementary Fig. S3F and G). Considering that DNA damage also leads to stalled cell cycle progression in the S phase(47,48), we examined whether the presence of glutamine affects cell cycle progression. With lower glutamine concentrations, more cells seemed to stay in the S phase and could not enter the G2/M phase, which was further aggravated by olaparib treatment, possibly due to unrepaired DNA damage (Fig. 3K and L). These results suggest that extracellular glutamine uptake by tumor cells is essential for promoting cell survival during PARPi and other DNA damaging-based chemotherapies.

### Blocking Microenvironment Glutamine Uptake Sensitizes Tumor Cells to Olaparib Therapy in Bone Metastasis

Our previous results suggest that extracellular glutamine is essential for tumor cell survival during PARPi therapy *in vitro* (Fig. 3E-H). To examine whether this is true *in vivo*, we knocked down the glutamine transporter SLC1A5 using short hairpin RNAs in SCP28 cells (Fig. 4A and B). SLC1A5 knockdown sensitized tumor cells to olaparib or cisplatin treatment, and the expression level of γH2AX increased dramatically, suggesting that glutamine uptake is essential for chromosomal stability in tumor cells (Fig. 4B and C). Consistently, SLC1A5 depletion also led to a significantly decreased GSH/GSSG ratio when cells were treated with olaparib, further confirming the importance of glutamine uptake in maintaining the redox balance in tumor cells (Fig. 4D). SCP28 cells with either vector control shRNA or SLC1A5-KD were then intracardially (IC) injected into 4-6 weeks old female nude mice. Ten days post-injection, mice were treated with either vehicle control or olaparib continuously until the experimental endpoint (Fig. 4E). As determined by BLI, KD of SLC1A5 or olaparib treatment reduced the bone metastasis burden. The strongest inhibitory effect was observed when olaparib was administered to SLC1A5-KD cells (Fig. 4F and G). Micro-computed tomography (μCT) imaging revealed that the KD of SLC1A5 reduced bone degradation. Similarly, olaparib treatment of the vector control cells also reduced bone degradation. The KD of SLC1A5 and olaparib treatment synergistically enhanced the bone-protection effect, leading to an almost intact bone structure. H&E and TRAP staining also revealed that olaparib treatment in SLC1A5 knockdown cells synergistically reduced the bone metastasis areas and the number of osteoclast cells associated with bone metastasis (Fig. 4G). Taken together, these *in vivo* experiments confirmed that blocking extracellular glutamine uptake effectively inhibits the progression of bone metastasis. Thus, our results demonstrate the importance of glutamine derived from the tumor microenvironment in generating resistance to PARPi therapy in bone metastasis.

**Figure. 4.**
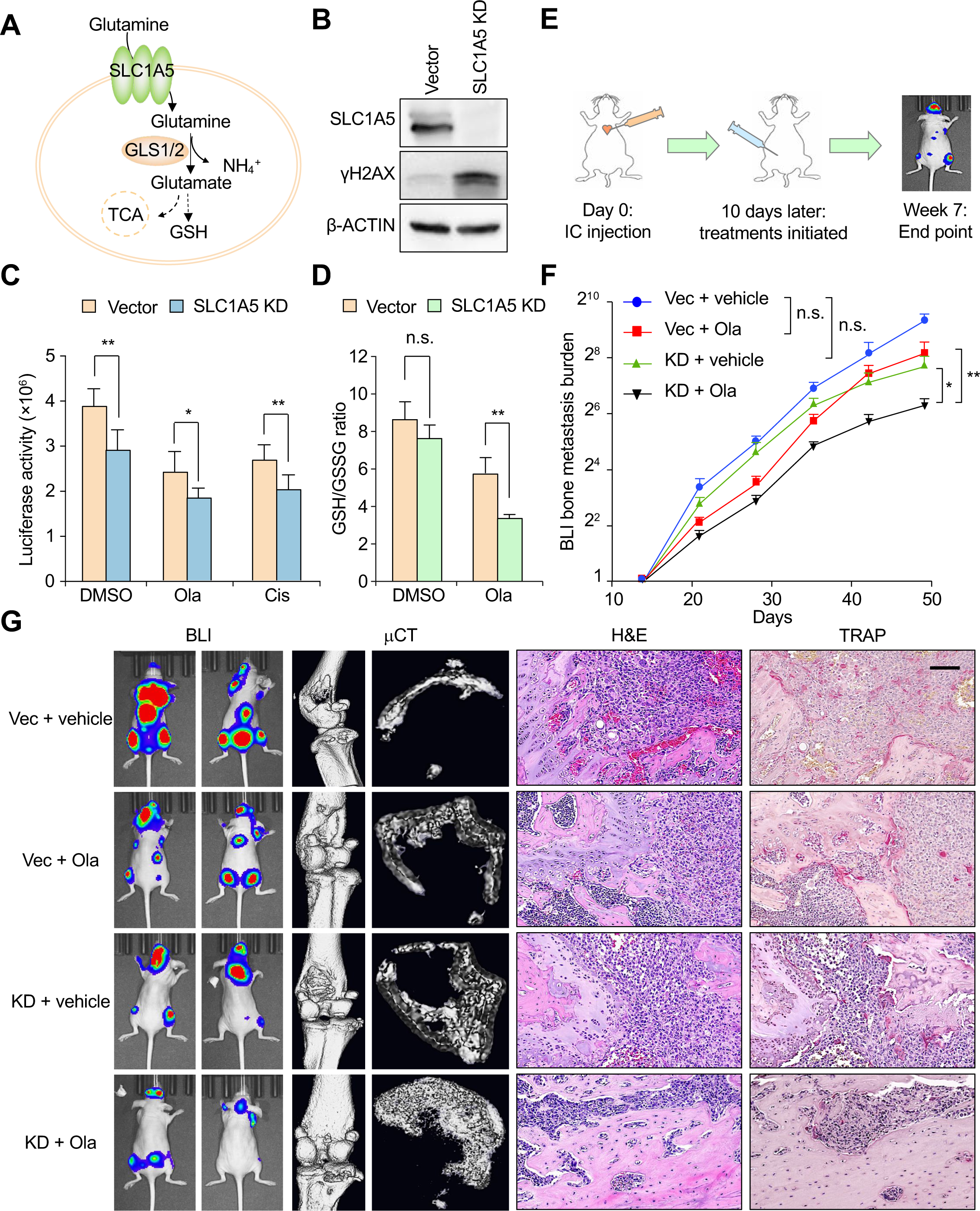
Inhibition of extracellular glutamine uptake sensitizes bone metastatic tumor cell to olaparib treatment *in vivo*. (**A**) A simplified model of the role of SLC1A5 in glutamine metabolism pathway. KD of SLC1A5 blocks the uptake of glutamine from tumor microenvironment and thus suppresses glutamine metabolism in tumor cells. (**B**) The expression level of γH2AX protein in SCP28-vector control or in SCP28-SLC1A5 KD cells was determined by immunoblotting. β-ACTIN was used as internal loading control. (**C**) Survival rates of SCP28-vector control or SCP28-SLC1A5 KD cells treated with either DMSO, olaparib (25 μM) or cisplatin (3 μM). Survival rates were measured as luciferase assay. (**D**) The ratio of GSH/GSSG in SCP28-vector or SCP28-SLC1A5 KD cells when treated with either DMSO or olaparib (25 μM). (**E**) Schematic illustration of the experimental procedure. 1×10^5^ SCP28 cells were IC injected into 4-6 weeks old female nude mice. Treatments were initiated 10 days post injection and continued until experimental endpoint. Mice were treated with either vehicle control or olaparib at 50 mg/kg with dosing schedule of 5 days on, 2 days off. (**F**) Quantification of bone metastasis burden as determined by weekly BLI. n = 12 mice per group. (**G**) Representative BLI images of mice in Week 7 post injection, μCT, H&E, and TRAP staining images at the/ experiment endpoint from experiment in E. Scale bar: 100 μm for H&E and TRAP. Data represents mean ± SEM. “*n.s.”* means not significant. *p < 0.05, **p < 0.01 by Student’s t-test (C, D, and F). **See also Supplementary Figure. S4**.

A previous report has suggested that PARPi may promote osteoclastogenesis. *In vitro* treatment with PARPi was proposed to induce osteoclast maturation (15). However, we noticed that low concentrations of PARPi were utilized *in vitro* with no direct evaluation of PARPi on osteoclastogenesis *in vivo*. Thus, a rigorous re-evaluation of the effect of PARPi on osteoclastogenesis, both *in vitro* and *in vivo* is required. We first tested the effect of olaparib on osteoclast maturation *in vitro* following a protocol similar to that described in Fig. 1A. When 20 µM olaparib was used, there was a significant decrease in the number of TRAP-positive osteoclasts (Supplementary Fig. S4A). To further confirm this effect *in vivo*, we treated 4-6 weeks old female nude mice with olaparib continuously for four weeks, and then evaluated bone integrity and the number of osteoclasts in the hind limbs of these mice. The bone tissues showed no obvious defects after olaparib treatment, as demonstrated by μCT imaging (Supplementary Fig. S4B). A detailed examination of TRAP-positive osteoclast cells in the hindlimbs revealed no increase but a mild decrease in the number of osteoclasts in PARPi-treated mice (Supplementary Fig. S4B and C). Based on this evidence, we conclude that PARPi does not induce osteoclast maturation. Instead, it may have slightly inhibited osteoclastogenesis *in vivo*.

### Combination Treatment of Zoledronate and Olaparib Synergistically Inhibits Bone Metastasis Progression

Since glutamine metabolism was significantly enhanced in mature osteoclasts, we examined the level of secreted glutamine in osteoclast CM using mass spectrometry. Macrophages or *in vitro* differentiated osteoclasts were cultured with glutamine-free culture media for 6 hours before CM collection. This protocol ensured the removal of any interference by glutamine present in the regular culture medium. Indeed, the concentration of glutamine in the CM from osteoclasts was significantly higher than that in the CM from macrophages (Fig. 5A). Thus, the higher glutamine concentration in osteoclast CM was because of the production and secretion from osteoclasts. To directly examine whether osteoclasts are the major stromal cells that provide glutamine to tumor cells, we followed the experimental procedure described in Fig. 5B. 6-8-week-old female nude mice were IC injected with SCP28 cells to induce bone metastasis. Mice were treated with either PBS or zoledronate. At the experimental endpoint, the mice were sacrificed, and the corresponding metastatic bone samples from the femur (red-dotted area) were minced for metabolomic analysis (Fig. 5B and Supplementary Fig. S5A). Although glutamine was present in significant amounts in PBS-treated bone samples, it decreased dramatically in the zoledronate-treated samples. The major downstream metabolites of the glutamine cytosolic pathway, including GSH, GSSG, and thus the total glutathione, were significantly downregulated in zoledronate-treated samples (Fig. 5C-F). Interestingly, the levels of another reducing power-providing metabolite, nicotinamide adenine dinucleotide (NAD), did not change after zoledronate treatment (Supplementary Fig. S5B). To eliminate any potential effects of varying tumor volumes on glutamine concentration in the previous experiment, we directly measured glutamine concentration in normal mouse bones treated with either PBS or zoledronate. The results indicated that both glutamine and GSH concentrations were reduced in zoledronate-treated normal mouse bones (Supplementary Fig. S5E and F). In summary, these results demonstrated that glutamine is enriched in osteolytic bone metastasis, which is likely derived from hyperactivated osteoclasts.

**Figure. 5.**
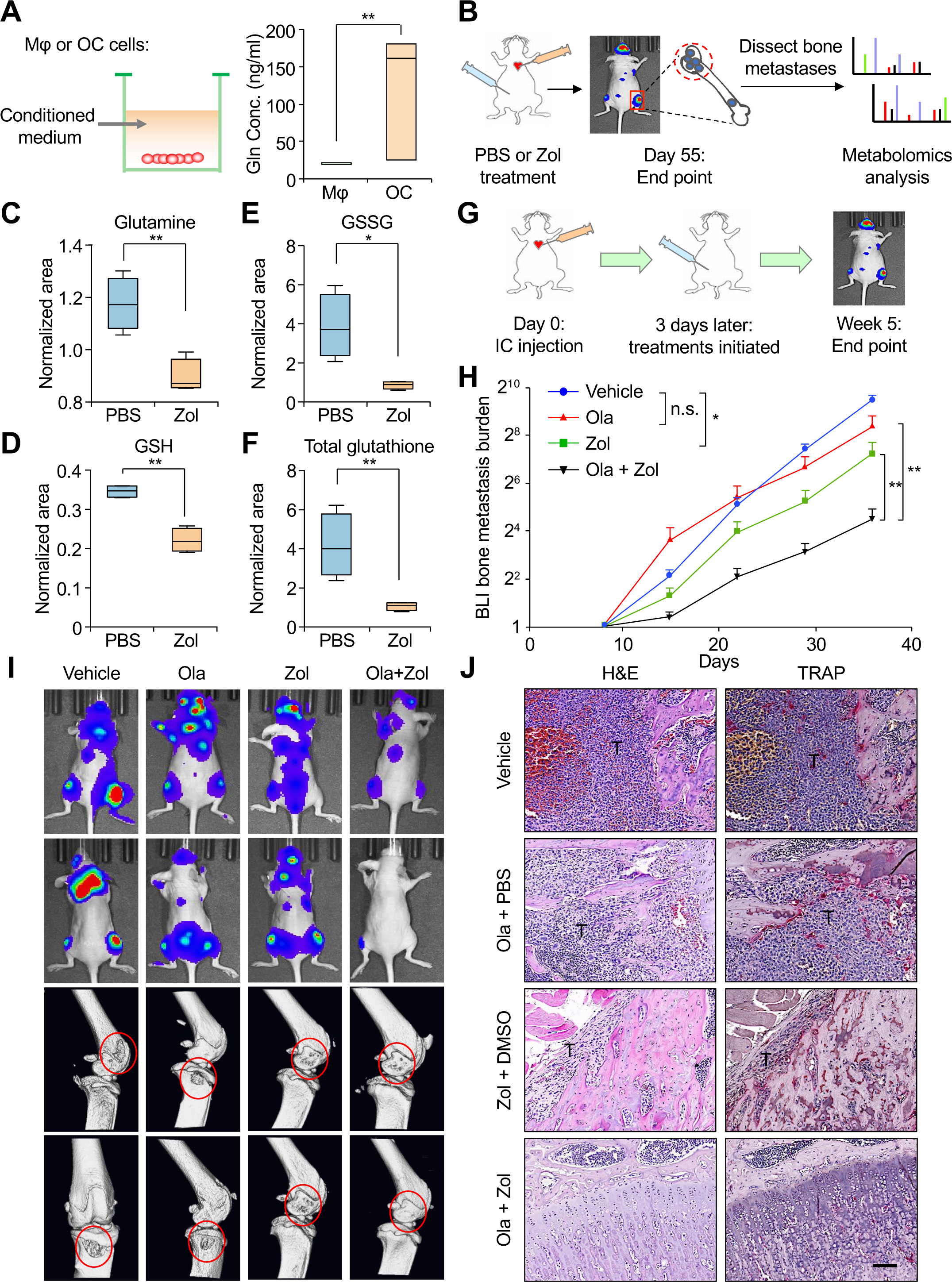
Combination therapy of zoledronate and olaparib synergistically inhibits bone metastasis progression. (**A**) The extracellular glutamine concentration of conditioned medium from either macrophages or from mature osteoclasts. Macrophages or *in vitro* differentiated osteoclasts were cultured with glutamine-free culture media for 6 hours before CM collection. The glutamine concentration in CM was determined by mass spectrometry analysis. (**B**) The workflow of analyzing metabolites in glutamine pathway in bone metastasis microenvironment. 1×10^5^ SCP28 cells were IC injected into 4-6 weeks old female nude mice. Treatment was initiated 10 days post injection and continued until experimental endpoint. Mice were treated with either vehicle control (PBS) or zoledronate (2 mg/kg) with the dosing schedule of twice a week. Mice were treated continuously for about 7 weeks. Bone metastasis was confirmed by BLI imaging and the corresponding metastatic bone areas (red-dotted area) were minced directly for sample collection and mass spectrometry analysis. (**C-F**) Experiment was performed as illustrated in B. Normalized areas of glutamine (C), GSH (D), GSSG (E), and total glutathione (F) in PBS-treated group or in zoledronate-treated group were presented. n = 4 per group. (**G**) Schematic illustration of the experimental procedure. 1×10^5^ SCP28 cells were IC injected into 4-6 weeks old female nude mice. Treatment was initiated 3 days post injection and continued until experimental endpoint. Mice were treated with either vehicle control, zoledronate, olaparib, or with combinatory treatment of both zoledronate and olaparib. The dosing schedule for olaparib (50 mg/kg) was 5 days on, 2 days off; while the dosing schedule for zoledronate (2mg/kg) was twice a week. (**H**) Quantification of bone metastasis burden as determined by weekly BLI. n = 12 mice per group. (**I**) Representative BLI and μCT images at experimental endpoint. Red-dotted circles indicate the osteolytic regions of bone metastasis. Zol represents zoledronate. (**J**) Representative H&E and TRAP staining images from experiment in **h**. Scale bar, 100 μm. Data represents mean ± SEM. “*n.s.”* means not significant, *p < 0.05, **p < 0.01 by Student’s t-test (A, C, D, E, F, and H). **See also Supplementary Figure. S5.**

Zoledronate is clinically approved for the treatment of bone metastasis in breast cancer. By inhibiting osteoclast function, zoledronate delays the progression of bone metastases and protects the bone from degradation, thus reducing bone-related adverse events in patients(49,50). As treatment with zoledronate drastically reduced the level of glutamine within the bone metastatic microenvironment, it may synergize with PARPi to inhibit bone metastasis. To test this hypothesis, we performed IC injection of SCP28 cells into 6 weeks old female nude mice and initiated treatment three days post injection (Fig. 5G). Treatment with olaparib slightly decreased bone metastatic burden by approximately 2.1-fold. Zoledronate inhibited bone metastasis burden by approximately 3.8-fold. Strikingly, when mice were treated with both olaparib and zoledronate, a 35-fold reduction in bone metastasis burden was observed (Fig. 5H and I). Hindlimb bones were analyzed using μCT, H&E, and TRAP staining. Compared to the vehicle control group, treatment with olaparib had no obvious effect on bone density or the number of TRAP-positive osteoclasts. Similar to previous reports, zoledronate inhibited osteoclastogenesis and increased the bone density. Moreover, combination therapy with PARPi and zoledronate generated a strong synergistic effect and significantly reduced the bone metastatic area, inhibited osteoclastogenesis, and protected bone integrity (Fig. 5I and J, Supplementary Fig. S5C and D, and Supplementary Videos S1-4). Thus, our results support the notion that blocking osteoclast activity with zoledronate synergistically suppresses the progression of bone metastasis following PARPi treatment.

### GSH, A Major Glutamine Downstream Metabolite, Neutralizes ROS during PARPi Therapy

Glutamine is utilized in two major metabolic pathways. Extracellular glutamine is taken up by the cells and metabolized into glutamate. Part of glutamate is then dehydrogenated into α-KG and enters the TCA cycle, which provides energy and carbon skeletons for cells. This part of glutamine metabolism is known as mitochondrial glutamine metabolism. The remaining glutamate is synthesized to GSH, the main reductive agent in cells, to resist oxidative stress (cytosolic glutamine metabolism) (51). Treatment with olaparib increased ROS levels in tumor cells, whereas higher concentrations of glutamine helped to reduce ROS levels (Fig. 6A and Supplementary Fig. S6A and B). We measured the intracellular concentrations of GSSG and GSH after olaparib or cisplatin treatment. Indeed, there were significantly increased concentrations of both GSSG and GSH, leading to an increased pool of total glutathione (Supplementary Fig. S6C). We noticed that with higher glutamine concentrations, the GSH/GSSG ratio in tumor cells was significantly increased in control cells, yet this ratio decreased upon olaparib or cisplatin treatment, indicating an enhanced conversion rate of GSH to GSSG, which is used to neutralize ROS caused by DNA-damaging agents (Supplementary Fig. S6C). Consistent with these data, we detected strong ROS induction with both olaparib and cisplatin treatment, but not with paclitaxel treatment (Supplementary Fig. S6D). This is likely the reason why CM from osteoclasts enhanced tumor cell survival after olaparib and cisplatin treatments, but not paclitaxel treatment (Fig. 1D and Supplementary Fig. S1B). Utilizing mitochondrial-localized catalase (mCAT), which protects cells from ROS(23), reduced tumor death induced by olaparib and cisplatin treatments (Supplementary Fig. S6E). These results highlight the essential role of ROS in the induction of cell death.

**Figure. 6.**
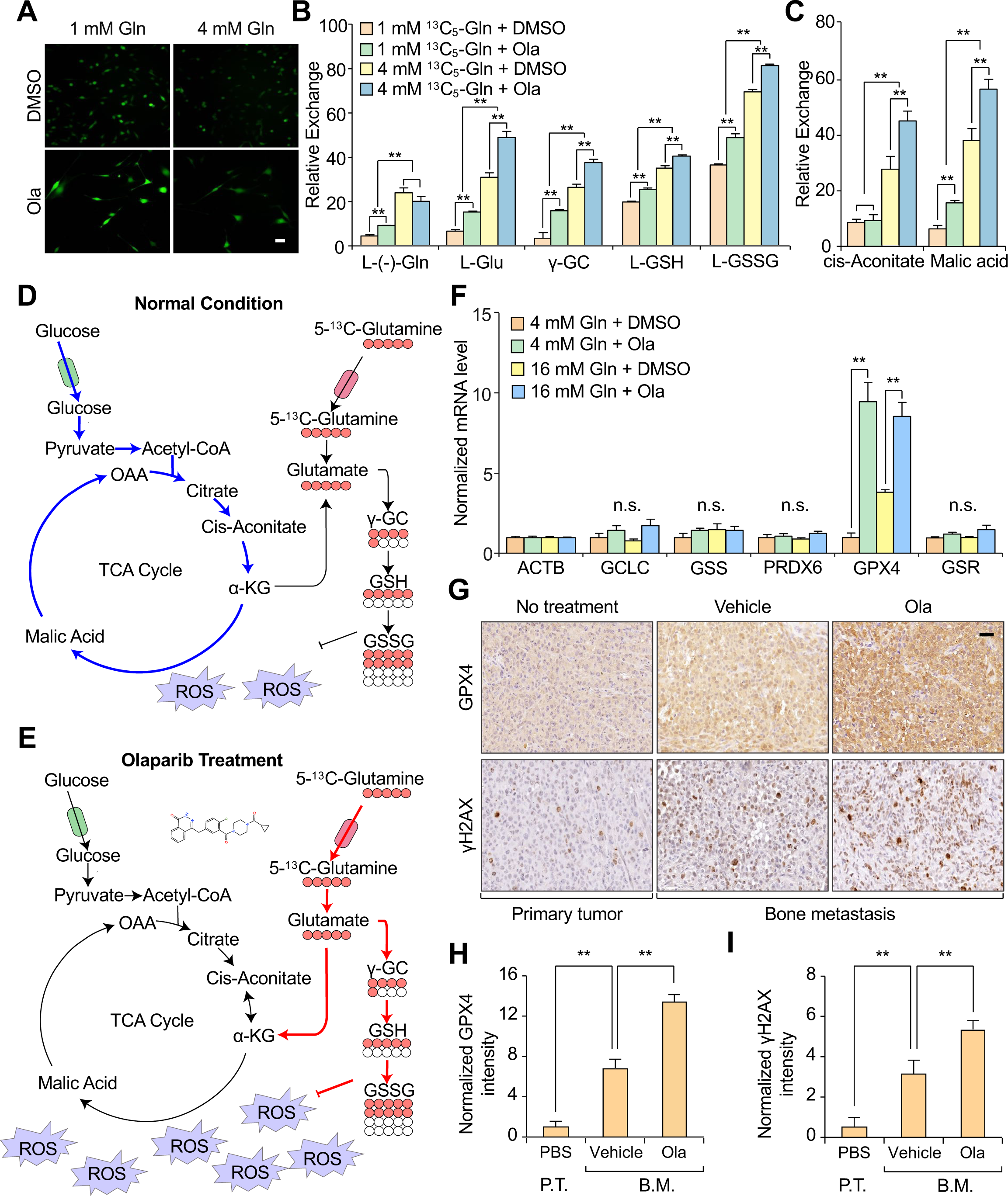
Glutathione production is enhanced in cancer cells to relieve oxidative stress during PARPi therapy. (**A**) SCP28 cells were treated with DMSO or olaparib in either low (1 mM) or normal (4 mM) glutamine media. The ROS level in these cells were determined by a 2’-7’dichlorofluorescin diacetate (DCFH-DA) ROS assay kit. Scale bar, 100 μm. (**B**) Metabolic flux analysis of relative levels of ^13^C-labeled major metabolites in cytosolic glutamine metabolism pathway in 4T1.2 cells. 4T1.2 cells were treated with either DMSO or olaparib (25 μM) in the culture media containing either 1 mM or 4 mM ^13^C-labeled glutamine. Cells were then lysed for mass spectrometry analysis. (**C**) Same experiment as in B. Metabolic flux analysis of relative levels of ^13^C-labeled major metabolites in mitochondria glutamine pathway (cis-Aconitate and Malic acid). (**D**) Schematic illustration of TCA cycle and glutathione metabolism in tumor cells treated with DMSO. Without olaparib, low level of endogenous ROS only requires baseline GSH and GSSG production in tumor cells. (**E**) Schematic illustration of TCA cycle and glutathione metabolism in tumor cells treated with olaparib. With olaparib induced DNA damage and ROS production, extracellular glutamine is then shunted to GSH and GSSG production to neutralize the increased ROS in cancer cells. Please note that in both D and E, the font size of each line reflects the activation status of each metabolic pathway. (**F**) SCP28 cells were treated with either DMSO or olaparib in the presence of 4 mM or 16 mM glutamine. The mRNA expression levels of genes encoding key enzymes in glutamine metabolism pathway were determined by qPCR. *ACTB* was utilized as internal loading control. n = 3 per group. (**G**) For primary tumor formation, SCP28 cells were mammary fat pad injected into 4-6 weeks old female nude mice. For bone metastases, SCP28 cells were IC injected into 4-6 weeks old female nude mice. Mice bearing bone metastases were treated with either vehicle control or olaparib. These tumor samples were collected at experimental endpoint for paraffin-fixing and then stained with antibodies against GPX4 protein or γH2AX. Scale bar, 40 μm. (**H-I**) Normalized IHC staining intensities of GPX4 (H) and γH2AX (I) from experiment in G using the ImageJ software. Data represents mean ± SEM. “n.s.” means not significant. **p < 0.01 by Student’s t-test (B, C, F, H, and I). **See also Supplementary Figure. S6.**

To systematically analyze the metabolic shunting of glutamine during olaparib treatment, we performed ^13^C isotope-labelled glutamine metabolism tracing experiments. Consistent with the results shown in Supplementary Fig. S6C, with a low glutamine concentration (1 mM), the uptake of extracellular glutamine, reduced L-glutathione, and oxidized L-glutathione increased dramatically after PARPi therapy (Fig. 6B). At the same time, mitochondrial glutamine metabolism was only mildly upregulated, as measured by the relative concentrations of cis-aconitic acid and malic acid, both of which are intermediates of the TCA cycle (Fig. 6C). In contrast, with sufficient glutamine supply (4 mM), both cytosolic glutamine metabolism and mitochondrial glutamine metabolism pathways were upregulated dramatically when cells were treated with PARPi (Fig. 6B and C). Thus, our results revealed a critical mechanism of glutamine metabolism. With limited glutamine available to tumor cells, glutamine is mostly directed to the cytosolic glutamine pathway to generate GSH as a reducing agent to neutralize endogenous ROS. However, when sufficient glutamine is available to tumor cells, it is utilized in both the TCA cycle and glutathione (GSH and GSSG) metabolism. In this situation, glutamine is not only utilized to generate GSH to neutralize DNA-damaging agent-induced ROS but is also utilized in the TCA cycle to provide energy and carbon resources to support cell proliferation (Fig. 6D and E).

SIRT4 has been reported to have a tumor-suppressive role and regulate the cellular metabolic response to DNA damage by inhibiting mitochondrial glutamine metabolism(52). To test whether olaparib treatment also influences mitochondrial glutamine metabolism by regulating the expression of SIRT family genes, we tested the mRNA expression levels of SIRT1-4. As a positive control, we treated the cells with camptothecin (CPT, a topoisomerase 1 inhibitor) and etoposide (ETS, a topoisomerase 2 inhibitor). A significant upregulation of SIRT4 mRNA was observed in positive controls but not in olaparib-or cisplatin-treated cells (Supplementary Fig. S6F and G). This result suggests that SIRT4 may not play a role in the resistance to PARPi therapy.

To understand why cytosolic glutamine metabolism, especially the conversion of GSH to GSSG, was strongly induced, we examined the mRNA expression levels of critical enzymes involved in glutamine metabolism (Supplementary Fig. S6H). Although the expression levels of most enzymes did not change, Glutathione peroxidase 4 (GPX4), an enzyme responsible for catalyzing GSH to GSSG, was significantly upregulated by olaparib treatment (Fig. 6F). We then examined the expression levels of GPX4 in primary tumor samples, vehicle-treated bone metastases, and PARPi-treated bone metastasis samples generated by SCP28 cell injection. A clear induction of GPX4 was observed in bone metastases compared to that in primary tumors, and the expression of GPX4 was further upregulated in PARPi-treated bone metastases.

Interestingly, the expression of γH2AX was also similarly regulated in these samples, suggesting that bone metastasis may induce much stronger oxidative stress and DNA damage than in the primary tumor, and that cytosolic glutamine metabolism is needed for bone metastasis (Fig. 6G-I).

### ATF4 Mediates the Induction of GPX4 Expression during PARPi Therapy

To directly examine the enriched metabolites in bone metastasis, we collected cancer cells from either primary sites or bone metastases and profiled their metabolomes (Supplementary Fig. S7A). Tumor cells that metastasized to the bone demonstrated clear metabolic reprogramming compared to the primary sites (Fig. 7A and Supplementary Fig. S7B and C). Through metabolic pathway enrichment analysis, glutathione metabolism was identified as the most enriched metabolic pathway in bone metastatic tumor cells (Fig. 7B). Importantly, both GSH and GSSG were significantly upregulated, whereas two other critical reducing power-providing metabolites, NAD and flavin adenine dinucleotide (FAD), did not change in bone metastases (Fig. 7C and Supplementary Fig. S7D). This is in agreement with data showing that bone metastatic tumor cells might experience a higher redox pressure than tumor cells at the primary site (Fig. 6G and H).

**Figure. 7.**
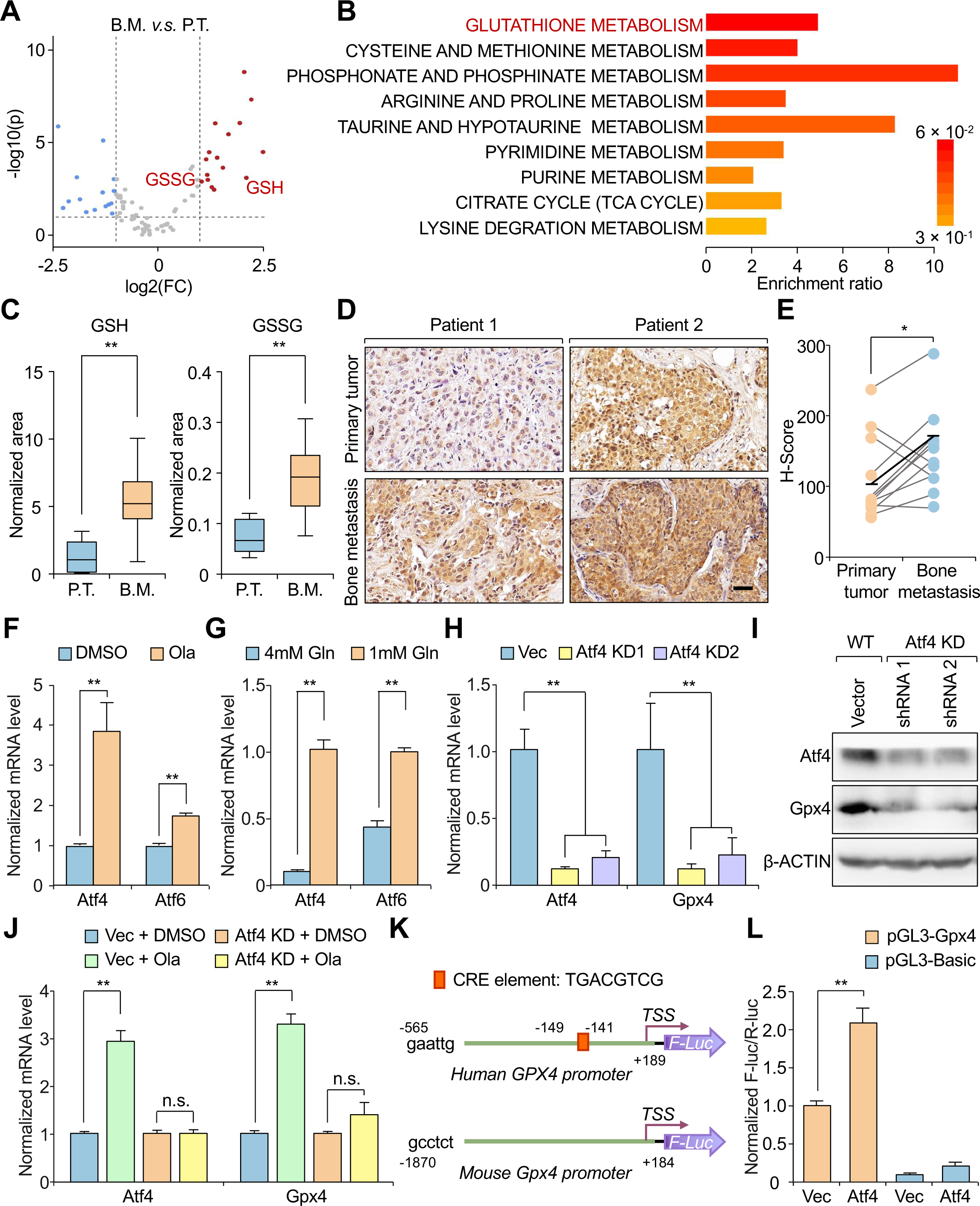
ATF4 mediates the induction of GPX4 expression during PARPi therapy. (**A**) Volcano plot of differentially expressed metabolites between bone metastatic samples (B.M.) and primary tumor samples (P.T.). Each dot represents one metabolite. Metabolites used in this analysis were changed for at least 1.75 folds and with false discovery rate (FDR) less than 0.05. (**B**) Metabolic pathway enrichment analysis of increased metabolites in bone metastatic samples *v.s.* primary tumor samples. (**C**) Normalized areas of GSH and GSSG in cancer cells from primary tumors or from bone metastases were presented. n = 7 for primary group; n = 14 for bone metastasis group. (**D**) Representative IHC staining images of GPX4 in paired human primary breast tumors and bone metastases. Scale bar, 100 μm. (**E**) IHC image scores (H-score, see METHODS for details) of GPX4 in paired primary tumors and bone metastases. Horizontal lines indicate median values. n = 11 per group. (**F**) mRNA expression levels of *Atf4* and *Atf6* was determined by qPCR in 4T1.2 cells treated with DMSO or 25 μM olaparib. *Actb* was utilized as internal loading control. n = 3 per group. (**G**) mRNA expression levels of *Atf4* and *Atf6* was determined by qPCR in 4T1.2 cells cultured in media containing 4 mM or 1 mM glutamine. *Actb* was utilized as internal loading control. n = 3 per group. (**H**) *Atf4* was KD by two independent shRNAs in 4T1.2 cells. mRNA expression levels of *Atf4* and *Gpx4* were determined by qPCR. *Actb* was utilized as internal loading control. n = 3 per group. (**I**) The protein expression levels of *Atf4* and *Gpx4* were determined by immunoblotting in vector cells and in *Atf4*-KD cells. β-actin was used as internal loading control. (**J**) 4T1.2-vector and 4T1.2-Atf4 KD (shRNA1) cells were treated with DMSO or 25 μM olaparib for two days. Cells were lysed for mRNA purification. mRNA expression levels of *Atf4* and *Gpx4* were determined by qPCR. *Actb* was utilized as internal loading control. n = 3 per group. (**K**) Schematic representation of *GPX4* promoter region used in reporter assay. The lengths of cloned promoters are indicated. Promoters used in the assay were −565 to +189 (human) and −1870 to +184 (mouse) relative to the transcriptional start sites. (**L**) 293T cells were transfected with indicated plasmids. Two days post transfection, Firefly luciferase activity was measured and normalized to Renilla luciferase control. Data represents mean ± SEM. *p < 0.05, **p < 0.01 by Student’s t-test (C, F, G, H, J, and L). **See also Supplementary Figure. S7.**

Based on these observations, we decided to use the expression of GPX4 as a surrogate marker for the activation of the glutathione synthesis (specifically for GSH to GSSG conversion). Eleven paired primary human breast tumor samples and corresponding bone metastatic samples were collected and immunohistochemically stained (IHC) for the GPX4 protein. Indeed, the expression level of GPX4 was significantly higher in bone metastases than in paired primary tumors (Fig. 7D and E).

The integrated stress response (ISR) is an evolutionarily conserved pathway that promotes cellular adaptation to diverse stresses and restores homeostasis(53–55). ATF4, the best-characterized effectors of ISR(56), belong to the activating transcription factor/cyclic AMP response element-binding protein (ATF/CREB family) (57–59). Recent reports indicate that ATF4 and its family member ATF6 are the major stress-induced transcription factors (TFs) involved in nutrient starvation, ROS stress, and the DNA damaging response(38,60,61), ATF4 and ATF6 may be responsible for GPX4 induction during PARPi therapy. Compared with the control group, PARPi treatment led to a more than 3-fold induction of *Atf4* expression and only a mild induction of *Atf6* expression (Fig. 7F). Interestingly, the expression levels of both *Atf4* and *Atf6* were induced in the culture media with low glutamine levels (Fig. 7G). The expression levels of *Atf4*, but not *Atf6*, also increased after cisplatin treatment (Supplementary Fig. S7E). Thus, we tested whether ATF4 was the major TF involved in GPX4 regulation. Using two independent shRNAs, the KD of *Atf4* significantly reduced the mRNA and protein levels of Gpx4 (Fig. 7H and I). Importantly, olaparib treatment induced *Gpx4* expression, which was completely abolished after *Atf4* KD (Fig. 7J). Based on these results, we cloned both the mouse and human *GPX4* gene promoters and performed a luciferase promoter assay. The expression of ATF4 significantly enhanced *GPX4* promoter activity (Fig. 7K and L and Supplementary Fig. S7F). These results suggested that the ATF4-GPX4 axis and the oxidation of GSH to GSSG are essential for promoting cell survival during PARPi therapy.

Our current results suggest that to cope with PARPi-induced ROS stress, bone metastatic cancer cells utilize osteoclast-derived glutamine, which are utilized for GSH synthesis. Simultaneously, PARPi treatment-induced stress increased ATF4 expression, which enhanced the mRNA transcription of *GPX4* genes. Whereas GPX4 catalyzes GSH into its oxidized form (GSSG) to counteract ROS and support tumor cell survival during PARPi therapy. Blocking osteoclast activity by zoledronate (or similar agents) suppressed microenvironmental glutamine supply, and synergistically eliminated bone metastatic tumor cells with PARPi therapy (Fig. 8).

**Figure. 8.**
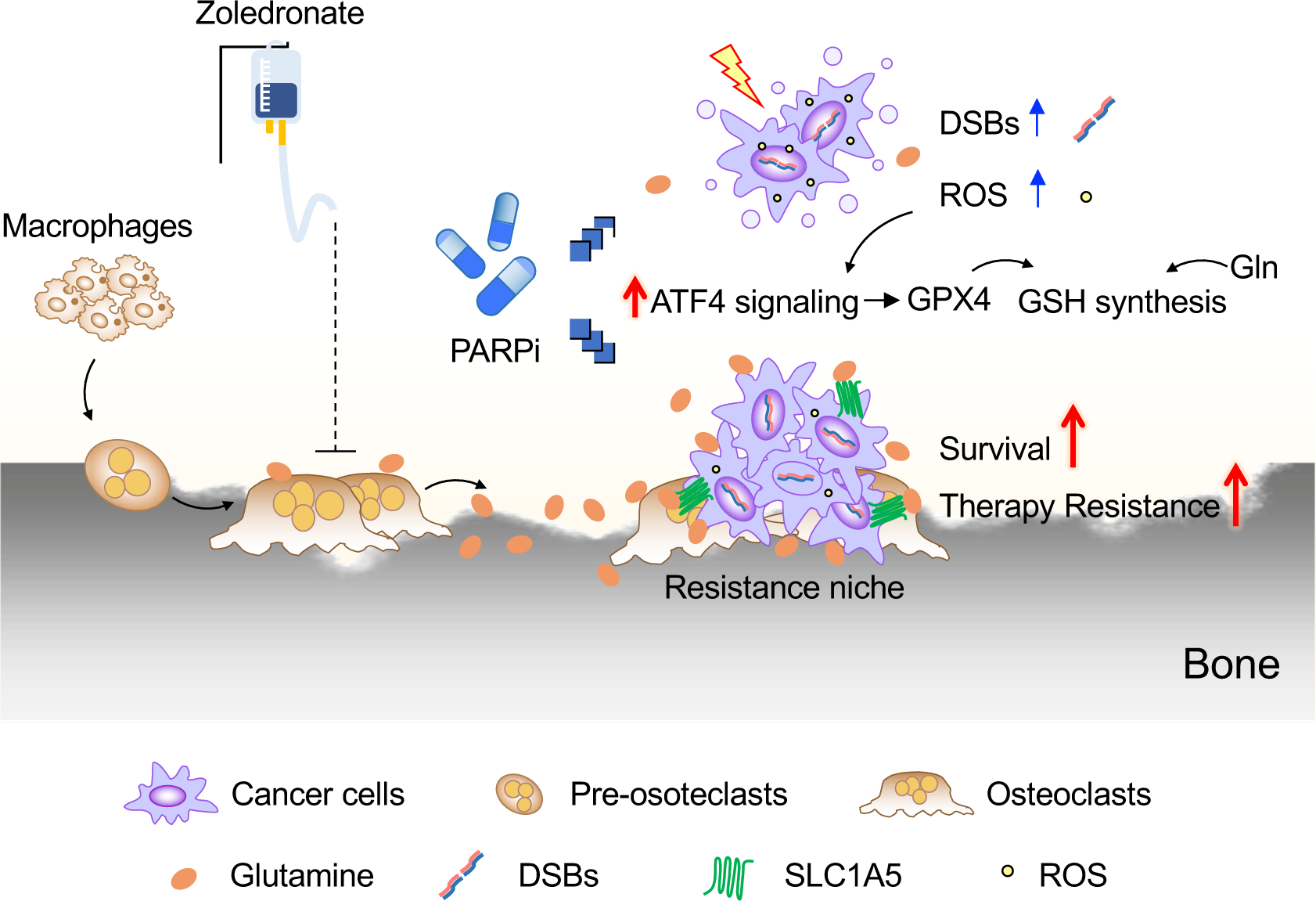
Schematic illustration of metabolic symbiosis between bone stromal and metastatic cancer cells in promoting therapy resistance. In osteolytic bone metastasis, hyper-activated osteoclasts provide increased supply of glutamine to bone microenvironment. On the other hand, PARPi treatment generates excessive ROS and DNA damage within cancer cells, which leads to enhanced GPX4 expression through ATF4 mediated transcriptional regulation. Collectively, increased environmental supply of glutamine and heightened enzyme activity of GPX4 accelerate glutathione metabolism to neutralize PARPi treatment-induced ROS. This leads to decreased level of DNA damage and enhanced cancer cell survival. Blocking osteoclast activity by zoledronate (or similar agents) could suppress tumor microenvironmental glutamine supply, and synergistically cause bone metastatic cancer cell death with PARPi therapy.3

## DISCUSSION

In this study, we conducted a metabolomic analysis of the metabolic pathways involved in osteoclastogenesis and identified several pathways that were upregulated, including the pathways that produce raw materials for GSH synthesis. We also found that the metabolic pathways in tumor cells shifted towards GSH metabolism during bone metastasis and DNA-damaging therapies. The increased microenvironmental amino acid supply, enhanced transportation through the cellular membrane, and accelerated GSH-to-GSSG conversion rate provide vital reducing power to support tumor cell survival during DNA-damaging therapies. Our findings demonstrate an orchestrated metabolic reprogramming of both bone osteoclasts and metastatic tumor cells that leads to the therapy resistance.

ROS play a complex role in cancer progression, with evidence suggesting that they may promote tumor initiation while also suppressing metastasis. Blocking intra-tumoral ROS to restore the redox balance in tumor cells is a common strategy to prevent tumor initiation(62,63). However, an increasing number of studies support that the increased ROS levels may suppress metastasis. Tumor cells can circulate and arrive at almost all distal organs, yet no overt metastasis can be established in skeletal muscle tissues due to dramatically increased ROS in this specific niche(23). Neutralizing ROS in tumor cells through the GSH precursor N-acetylcysteine increased the metastatic burden in a melanoma model. Likewise, genetic or pharmacological blocking of GSH synthesis in colon cancer cells suppressed liver metastasis(22). Bone is a common metastatic site for breast cancer and other cancer types(64)(*65*), yet whether and how tumor cells balance the enhanced ROS during bone metastasis progression has not been investigated. By profiling the metabolome of osteoclasts, we found that glutamine level was significantly upregulated during osteoclastogenesis, this is in consistent with a previous report that glutamine production is likely enhanced through likely through the utilization of branched-chain amino acids (BCAAs)(65). Thus, hyperactivation of osteoclasts provides key metabolic support for tumor cells to synthesize GSH and maintain redox balance. In turn, tumor cells upregulate expression of GPX4 responsible for GSH-to-GSSG conversion, to enhance GSH production and oxidation, which provides essential reducing power to neutralize ROS. This metabolic symbiosis between osteoclasts and tumor cells provides another layer of tumor-stromal communication to promote bone metastasis and therapy resistance.

Treatment with PARPi or other DNA-damaging agents significantly increased the levels of damaged DNA and cellular ROS. The integrated stress response (ISR) is reported to be activated during such stress condition(53–55). Our study confirmed that ATF4, the main transcription factor in the ISR(38,60,61), is significantly induced in cancer cells during treatment with PARPi and cisplatin. In addition, we found that ATF4 regulates the transcriptional activation GPX4. Therefore, our study identified an ATF4-GPX4 axis that promotes the GSH/GSSG metabolism pathway in resistance to bone metastasis therapy. Since ATF4 is the core TF in ISR, it is thus possible that other stress-induced survival signaling or metabolic pathways are activated in tumor cells, which would be worth investigating in the future.

Glutamine is crucial for the synthesis of GSH. Metastatic cancer cells experience increased ROS stress at distal sites(23), the ROS stress is further amplified through DNA-damaging therapies(24,25). Thus, these cancer cells are in exigent need to synthesize GSH and to neutralize ROS. However, cancer cells are not able to *de novo* synthesis sufficient glutamine to support their own needs under these stressed conditions, and they rely on glutamine acquired from the microenvironment(66,67). For example, tumor cells are dependent on absorbing extracellular glutamine to support the proliferation(68)(71)(*72*). However, the stromal cell type that provides glutamine during bone metastasis has not been appreciated. Our results showed that osteoclasts produce much higher concentrations of glutamine than that from macrophages. Furthermore, bone tissues from mice treated with zoledronate, which suppresses osteoclast activity, showed a dramatic decrease of these two amino acids. These results strongly support the notion that osteoclasts are the major source of glutamine in bone. Targeting the supply of glutamine could be a potential strategy for generating synergistic effects with PARPi treatment. Indeed, combination therapy with PARPi and zoledronate, an osteoclast inhibitor, significantly reduced the bone metastasis burden and protected bone integrity compared with single treatment. We expect that denosumab, a monoclonal antibody that targets RANKL and osteoclastogenesis, will also synergize with PARPi to suppress bone metastasis. As both olaparib and zoledronate (or denosumab) are FDA-approved pharmaceuticals for the treatment of breast cancer patients, their clinical usage should be tested in the future.

## METHODS

## RESOURCE AVAILABILITY

### Lead contact

Further information and requests for resources and reagents should be directed to and will be fulfilled by lead contact Dr. Hanqiu Zheng.

### Materials availability

This study generated multiple knockdown constructs that were made available through standard Material Transfer Agreement (MTA).

### Data and code availability

RNA sequencing data are available at the following website (GEO Accession Number: GSE208234): https://www.ncbi.nlm.nih.gov/geo/query/acc.cgi?acc=GSE208234 (reviewer token: ypmdcowstjadfwp).

## EXPERIMENTAL MODEL AND SUBJECT DETAILS

### Animal studies

All procedures involving mice and experimental protocols were approved by the Institutional Animal Care and Use Committee (IACUC) of Tsinghua University. For orthotopic primary tumor formation, female nude mice (4-6 weeks old) were anesthetized and a small incision was made to reveal the mammary gland. 1×10^5^ SCP28 tumor cells suspended in 10 μl PBS were injected directly into the mammary fat pad. For experimental bone metastasis, an intracardiac (IC) injection was used. Tumor cells were harvested from subconfluent cell cultures, washed with PBS, and suspended at 1×10^6^ cells/ml in PBS. The mice were anesthetized before injection. For IC injection, 0.1 ml cells were injected into the left cardiac ventricle of 4-6-weeks-old, female BALB/c mice or athymic nude mice using 26G needles. Successful injection was confirmed by BLI imaging, with an evenly distributed bioluminescent signal throughout the body. Bone metastasis burden was monitored using weekly BLI and μCT imaging. At the experimental endpoint, the mice were euthanized for bone histology analysis and half of the bone samples were subjected to μCT imaging and bone density quantification. For *in vivo* inhibitor treatment, mice were treated with vehicle control or 50 mg/kg of olaparib with a dosing schedule of five days on, two days off. Zoledronate (Adamas Reagent, 161728) was administered to the mice at a dose of 2 mg/kg with dosing schedule of twice a week.

### Cell culture

HEK293T (CRL-11268), 4T1.2 (CRL-11268), and SCP28 and related knockdown cells were cultured in Dulbecco’s modified Eagle’s medium (DMEM, Gibco) supplemented with 10% heat-inactivated Fetal Bovine Serum (FBS, Gemini) and 1% penicillin/streptomycin (Corning). To generate stable knockdown cell lines, a pLKO.1-based lentiviral vector was used. Lentiviruses were packaged into HEK293T cells. Conditioned media from packaging cells containing viruses were collected 2 and 3 d after transfection. Recipient cell lines were exposed to conditioned media containing viruses supplemented with 8 μg/ml polybrene for 24 h. Infected cells were selected using puromycin to generate stable cell lines. HEK293T cells were obtained from human females. All the other cancer cell lines were breast cancer cell lines obtained from either female human or mice.

### Osteoclast differentiation assay and cancer cell killing assay with chemotherapeutic treatments

Macrophages were isolated from bone marrow cells flushed from the tibia of 6-week-old C57/BL6 mice and filtered through a 70 μm cell strainer before overnight culture in α-MEM with 10% FBS. The following day, non-adherent cells were plated and supplemented with 50 ng/ml M-CSF for 2 days. Cells (2×10^5^) were then re-plated in each well of 12-well plates in the presence of 100 ng/ml RANKL in DMEM containing 10% FBS, with the media changed every 2 days (media was changed twice a day from Day 5 and on). Matured osteoclasts were TRAP-stained using a TRAP staining kit (38A-1KT, Sigma), and TRAP-positive and multinucleated cells were quantified as mature osteoclasts. Macrophage or osteoclast CM was collected when mature osteoclasts were present in the culture plates. The CM was then centrifuged at 500 × g for 3 min to remove insoluble debris and then filtered through an Amcion Ultra-15 Centrifugal filter, 3 kDa (Catlog # UFC900324, Sigma). 4×10^4^ SCP28 or 4T1.2 cells were plated in each well of 24-well plates treated with either olaparib (25 μM), cisplatin (3 μM), vincristine (10 nM) or paclitaxel (25 nM) in the presence of macrophage CM or osteoclast CM mixed with regular culture medium at the ratio of 1:1.

### Metabolomics analysis

Macrophages or osteoclasts were extracted using ice-cold 80% methanol and subjected to three rapid freeze-thaw cycles. The cell lysate was centrifuged at 15,000 rpm for 15 min at 4 °C. The supernatant containing aqueous metabolites was evaporated to dryness using a concentrator. The metabolites were reconstituted in 50 μl of 0.03% formic acid in analytical-grade water, vortexed, and centrifuged to remove insoluble debris. The supernatant was transferred to an HPLC vial for metabolomic analysis. The peak area for each detected metabolite was normalized against the total ion current of all the detected metabolites within the sample to correct for any possible variations. The normalized area values were used as variables for statistical data analysis using the Student’s t-test.

### Bone histology analysis

Hindlimb bones were excised from mice at the end of each experiment immediately after the last BLI imaging. Tumor-bearing hind limb bones were fixed in 4% neutral-buffered formalin, decalcified in 10% EDTA (pH 7.4) for 2 weeks, and embedded in paraffin for hematoxylin and eosin (H&E) or tartrate-resistant acid phosphatase (TRAP) staining. Osteoclast number was assessed as multi-nucleated TRAP-positive cells and reported as the number per field.

### X-ray imaging and μCT analysis

Femurs and tibias were dissected from the mice and scanned using a Quantum GX (PerkinElmer). μCT images were acquired at 88 kV and 88 μA, X-ray images were acquired at 88 kV and 40 μA, and images were acquired using high-resolution model scanning for 4 min, resulting in a voxel size of 72 μm × 72 μm. Images were reconstructed using a Quantum GX system and analyzed using a Quantum GX viewer (PerkinElmer). Regions with significant osteolytic bone degradation were segmented manually. The bone degradation area was measured using Photoshop software.

### TRAP (tart-resistant acid phosphatase) staining

Sections of 4% formalin-fixed, ethylenediaminetetraacetic acid (EDTA)-decalcified, and paraffin-embedded bones (with osteoclasts) were deparaffinized. Slides were incubated in substrate incubation buffer (2.5 ml Naphthol-ether solution (20 mg naphthol AS-BI phosphate (Sigma, N-2125) in 2.5 ml ethylene glycol monoethyl ether (Sigma, E-2632)) mixed with 250 ml basic medium (3.8 g Sodium acetate·3H_2_O, 2.9 g L-(+) tartaric acid 0.7 ml in 250 ml distilled water)) at 37℃ for 30 min. The color reaction medium was prepared as (sodium nitrite solution (4% sodium nitrite (m/v)) 1:1 mixed with pararosaniline dye stock (1 g pararosaniline chloride (Sigma, P-3750) in 20 ml 2N HCl)) and allowed to stand for 2 min. The colored reaction medium was then mixed with 25 times the volume of basic medium. The slides were then incubated in this diluted color reaction medium until TRAP-positive osteoclasts were visible under a microscope. The slides were then rinsed in water and subjected to hematoxylin staining for 10 s.

### Isotype tracing analysis with ^13^C-labeled glutamine

1×10^6^ 4T1.2 cells were plated on 10 cm dishes and cultured in unlabeled glutamine-containing DMEM medium (Gibco) for 24 h, then switched to ^13^C labeled glutamine-containing medium (1 mM or 4 mM ^13^C_5_-glutamine) for 2 days with or without 25 μM olaparib treatment. To harvest intracellular metabolites, the medium was aspirated from the cell culture, trypsinized, and collected by centrifugation at 1000 rpm for 5 min. The pellets were washed twice with cold phosphate-buffered saline (PBS). Metabolites were quenched with 80% cold methanol and the resulting lysates were stored at −80 °C for 1 h. The cell lysates were then centrifuged at 4 °C to remove proteins, and the supernatant was collected and evaporated using a nitrogen evaporator. The processed samples were sent to the PKPD platform of Tsinghua University for ^13^C-isotope tracing to identify the essential metabolites in the glutamine pathway.

### Luciferase assay

For the luciferase assay, the cells were lysed using luciferase lysis buffer (2 mM EDTA, 20 mM DTT, 10% glycerol, 1% TritonX-100, 25 mM Tris base, pH 7.8, adjusted with H_3_PO_4_) for 1 h at room temperature. 25 μl lysate was added to each well of a 96 well white plate (3 repeats per sample), 75 μl luciferase substrate solution was added into each well, and luciferase activity was assayed using a Multiple Microplate Luminometer (EnVision).

### Generation of stable knockdown cell lines

To knock down endogenous GLS1, SLC1A5, or ATF4 in SCP28 and 4T1.2 cells, short hairpin RNA (shRNA) constructs targeting human or mouse GLS1, SLC1A5, or ATF4 were generated. Briefly, human or mouse *GLS1*, *SLC1A5*, or *ATF4* short hairpin RNA (shRNA) sequences were cloned into the pLKO.1 vector (Addgene_13425). The following shRNA sequences were used for human GLS1:5’-GCACAGACATGGTTGGTATAT-3’ and 5’-GCCCTGAAGCAGTTCGAAATA-3’; mouse GLS1:5’-GAGGGAAGGTTGCTGATTATA-3’ and 5’-ATCTCGACGGGTTGCTATAAT-3’; human SLC1A5:5’-CTGGATTATGAGGAATGGATA-3’; mouse ATF4:5’-CTAGGTCTCTTAGATGACTAT-3’ and 5’-CGGACAAAGATACCTTCGAGT-3’. Stable knockdown of these genes was achieved using lentiviral infection and selection with 3 μg/ml puromycin for 3-5 days and the knockdown efficiency was confirmed by immunoblotting.

## Reverse transcription and qPCR

Total RNAs was isolated from the cells following the manufacturer’s instructions. RNAs was reverse transcribed into cDNAs using a Reverse Transcription kit (Promega, A2790). Real-time quantitative PCR (qPCR) was performed using SYBR Green PCR Mix (DSBIO, P2105) on a Bio-Rad CFX96 PCR system. The gene-specific primer sets were used at a final concentration of 0.5 μM. All real-time q-PCR assays were performed in triplicate in at least two independent experiments. The relative expression of each target gene was normalized to *GAPDH* or *ACTB* mRNA levels.

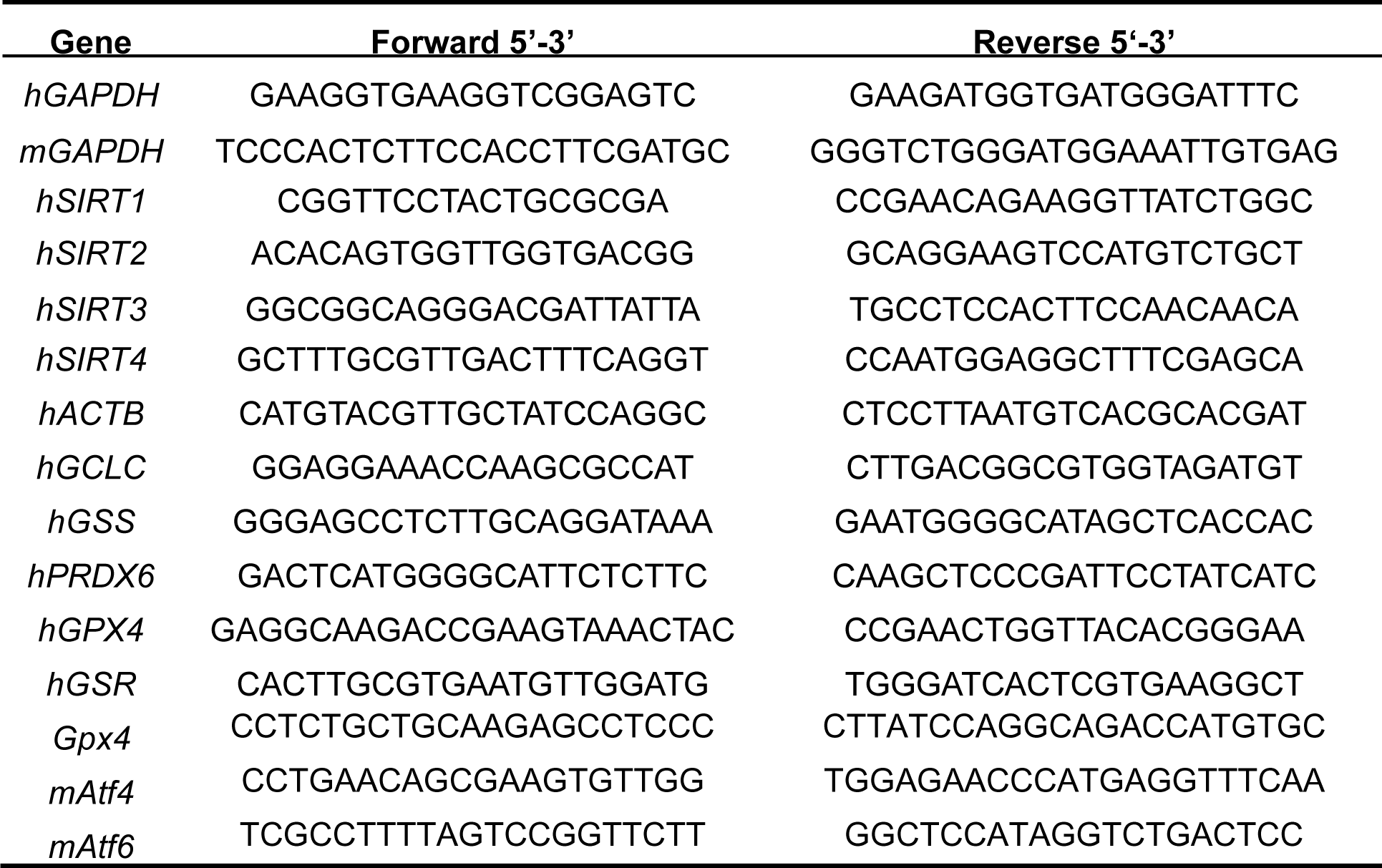

### GPX4 promoter luciferase assay

Around 800 bp human GPX4 promoter region (−565 to +189 bp relative to the transcriptional start site) or 2 kb mouse Gpx4 promoter region (−1870 to +184 bp relative to the transcriptional start site) was cloned from genomic DNAs from MDA-MB-231 cells or mouse liver and inserted into pGL3-Basic vector. The primers used for human GPX4 promoter cloning were CGCGGTACCGAATGGTGGATGAGCCTGTT (forward primer, KpnI site) and CGCCTCGAGCAAGGTCACTGGAGCCTGAG (reverse primer, XhoI site). The primers used to clone the mouse Gpx4 promoter were CGCGGTACCGCCTCTCAGCACTCTCAAAACAG (forward primer, KpnI site) and CGCCTCGAGATAGCCGGCCATTCCGAGGAG (reverse primer, XhoI site). The pLEX plasmids containing cDNAs for human ATF4 and mouse Atf4 were generated by PCR from the MDA-MB-231 or mouse liver cDNA library and inserted into the pLEX vector using Mul1 and Age1 restriction sites. The primer sequences used were CACACGCGTGCCACCatgaccgaaatgagcttcctg (forward) and CACACCGGTctaggggacccttttcttccccct (reverse) for human ATF4, and CACACGCGTGCCACCatgaccgagatgagcttcctgaac (forward) and CACACCGGTttacggaactctcttcttcccccttg (reverse) for mouse Atf4.

For the luciferase reporter assay, 2×10^5^ 293T cells were seeded in 12 well plates. Cells were transfected with the luciferase promoter constructs pMSCV-Renilla control, pLEX-hATF4, or pLEX-mATF4. Two days after transfection, the cells were lysed and luciferase activity was assayed using a Multiple Microplate Luminometer (EnVision). Renilla luciferase activity was used as an internal control.

### Immunofluorescence (IF) staining

The cells (1×10^5^) were seeded onto sterile glass coverslips in 12-well plates. The cells were treated with DMSO, olaparib (25 µM), cisplatin (3 µM), or V-9302 (9 µM) for about 2-3 days. Cells were washed with cold PBS, fixed with 4% formaldehyde for 10 min, permeabilized with 0.2% Triton X-100 for 5 min, blocked with 5% BSA in PBS for 1 h, incubated with appropriate primary antibodies overnight at 4°C, and then with secondary antibodies for 1 h at 4°C in the dark. The antibodies used for IF staining were anti-γH2AX (1:400; CST, 9718S), anti-Cleaved Caspase-3 (Asp175) (1:400; CST, 9661S). The secondary antibody used in the experiments was goat-anti-rabbit, Daylight 594 (1:100; Abbkine, ABB-A23420).

### GSH/GSSG and ROS measurements

To measure the GSH/GSSG ratio, 4T1.2 or SCP28 cells were seeded in 10 cm dishes at a density of 1×10^6^ cells per plate and cultured in regular DMEM. The medium was changed to DMEM containing 4 mM or 1 mM glutamine before treatment with DMSO or 25 μM olaparib. Two days later, the cells were collected, and intracellular GSH and GSSG concentrations were measured using a GSH/GSSG measurement kit (Beyotime, S0053) following the manufacturer’s protocol. To measure ROS, 4T1.2 or SCP28 cells were seeded in a 12-well plate at a density of 1-5×10^4^ cells per well. The culture medium was changed to DMEM containing 4 mM or 1 mM glutamine before treatment with DMSO or 25 μM olaparib. Two days later, the cells were stained using the ROS indicator DCFH-DA kit (Solarbio, D6470), following the manufacturer’s protocol.

### Cell cycle analysis

For cell cycle analysis, the cells were collected, fixed in 70% cold ethanol for 4 h, and treated with RNase A for 30 min. The cells were stained with propidium iodide (PI) for 30 min before FACS analysis.

### Colony formation assay

A total of 4×10^3^ cells were seeded per well in 6-well plates. The cell culture medium was replenished every three days. The colonies were cultured for two weeks before fixing, stained with Sulforhodamine B (SRB), and counted. For inhibitor treatment in the colony formation assay, cells were treated with DMSO, 25 μM olaparib, or 3 μM cisplatin.

### Immunoblotting analysis

For immunoblotting analysis, whole-cell lysate samples were collected using a cell lysis buffer (50 mM Tris-HCl pH 7.4, 150 mM NaCl, 1 mM EDTA, and 1% NP-40). Lysates were heated to denature the proteins, loaded onto SDS-PAGE gels for electrophoresis, and subsequently transferred to PVDF membranes (Millipore). Membranes were blocked in 5% milk for 1 h at room temperature prior to overnight incubation with primary antibodies. The primary antibodies used were anti-β-actin (1:5000, Abcam, ab6276), anti-GLS1 (1:1000, CST, 56750S), anti-GPX4 (1:2000, Abcam, ab125066), anti-ATF4 (1:1000, CST, 11815S), and anti-ASCT2 (SLC1A5) (1:1000, CST, 8057S). Membranes were incubated with horseradish peroxidase (HRP)-conjugated anti-mouse or rabbit secondary antibodies (1:10000, EASYbio, BE0102-100/BE0101-100) for 1 h at room temperature, and chemiluminescence signals were developed using an ECL substrate at a ratio of 1:1 (Tanon, 180-5001) on a Champchemi digital image acquisition machine (Sagecreation). Image quantification was performed using ImageJ software (NIH).

### Immunohistochemical (IHC) staining

For immunohistochemical staining, antigen retrieval with citric buffer (pH 6.0) was performed on the paraffin-embedded sections. The sections were incubated with primary antibodies overnight at 4 °C and with secondary antibodies (DaKo REAL EnVision, K5007, HRP Rabbit/Mouse) for 1 h at room temperature. DAB substrate kit (ZSGB-BIO, ZLI-9019) was used as the chromogen. Tissue sections were counterstained with hematoxylin (Sigma-Aldrich), dehydrated, and mounted. The primary antibodies used for immunohistochemical staining were anti-GPX4 (1:800, Abcam, ab125066) and anti-γH2AX (1:500, CST, 9718S).

### Metabolomics analysis of bone metastasis tumor tissues

Bone metastasis samples were collected from the mice that received weekly treatment with PBS or zoledronate at 2 mg/kg. Bone metastatic tissues were acquired using BLI. Freshly collected samples were preserved in 1.5 ml eppendorf tubes, immersed in liquid nitrogen to quench immediately, and frozen at −80 °C until metabolomic analysis was commenced. Each sample was added to 200 μl 80% methanol (including 1% PFA) and homogenized. An extraction solution (80% methanol with 1% PFA), 19 times the original sample weight, was added to the homogenized sample. After slow vortexing, the supernatant was transferred to a fresh Eppendorf tube. Quality control (QC) samples were prepared by mixing 20 μl of each extract. 100 μl sample was transferred into a new Eppendorf tube and mixed with 900 μl extraction solution. Each sample (900 μl of each sample was placed in a fresh 1.5 ml Eppendorf tube before drying in a vacuum concentrator for further LC/MS (Liquid Chromatography-Mass Spectrometry) analysis.

### Paired human primary breast tumor and bone metastasis sample analysis

Eleven paired primary breast tumor and bone metastasis samples were collected at the Department of Orthopedic Oncology, Affiliated Hospital of Qingdao University. The experimental procedures were approved by the Human Ethics Committee of the Affiliated Hospital of Qingdao University. Fresh specimens were collected at the time of surgical resection, under the supervision of qualified pathologists. Paraffin-embedded samples were processed and sequential tumor slides were generated for IHC staining.

### RNA-seq analysis

5×10^5^ MDA-MB-231 cells were plated on 10 cm dishes and cultured for 24 h. The cells were then treated with either DMSO or 40 µM olaparib for three days. Cells were lysed for RNA extraction, cDNA library construction, and RNA-seq analysis. Raw data were processed through quality control using FastQC(69)(72)(*74*). The sequencing data were aligned using STAR(70)(73)(*75*), SAMtools(71)(74)(*76*), and Subread(72)(75)(*77*). Downstream data analysis was performed using R program(73)(76)(*78*) and R packages, including edgeR(74) (77)(*79*) and limma(75)(78). Metabolic gene sets that were used to match metabolic genes to the top genes in the RNA-seq data were downloaded from MSigDB(35–37) and KEGG(https://www.kegg.jp/). Gene set enrichment analysis was performed using the Metaspace(76)(79).

## QUANTIFICATION AND STATISTICAL ANALYSIS

The results are reported as the mean ± standard deviation (SD) or mean ± standard error of the mean (SEM), as indicated in the Fig. legend. Statistical comparisons were performed using an unpaired two-sided Student’s t-test with unequal variance assumption. All experiments with representative images (including immunoblotting and immunofluorescence staining) were repeated at least twice and representative images are shown.

## ACKNOWLEDGEMENTS

We thank Z. Chang for helping with IHC staining, and Y. Kang, Y. Fu, P. Jiang for their helpful discussions. We thank C.J. David for proofreading this manuscript. We thank all the members of the Zheng Laboratory for their helpful discussions and technical assistance. We thank the Technology Center for Protein Sciences at Tsinghua University for mass-spectrometry support, the Laboratory Animal Research Center, and the Center of Biomedical Analysis at Tsinghua University for animal research support. This study was partially supported by the National Key Research and Development Program of China (2020YFA0509400 to H.Z.), National Science Foundation of China (81772981 and 81972462 to H.Z.), Tsinghua University Initiative Scientific Research Program, and the Tsinghua-Peking Center for Life Sciences.

## AUTHOR CONTRIBUTIONS

H.F. and Z.X. designed and performed the experiments and analyzed the data. Z.H. designed the study; H.F., Y.Z., and K.Y. performed the experiments for metabolic analysis and helped with data analysis. Z.X. performed the glutathione/ROS measurement assays and RNA-seq data analysis with the help of T.Z., X.W., H.H., and X.L. provided technical assistance for the animal studies. B. Z. and B.Y. collected and provided paired clinical breast cancer patient samples from primary breast cancer tumor tissues and bone metastatic tumor tissues. H.Z. developed the study concept, designed the experiments, and supervised the study. H.Z., H.F. and Z.X. wrote and revised the manuscript.

