## Supplemental Information for "Osteoclast-Cancer Cell Metabolic Symbiosis Renders PARP Inhibitor Therapy Resistance in Bone Metastasis"

**Supplemental Data**

Figure S1 is related to Figure 1.
Figure S2 is related to Figure 2.
Figure S3 is related to Figure 3.
Figure S4 is related to Figure 4.
Figure S5 is related to Figure 5.
Figure S6 is related to Figure 6.
Figure S7 is related to Figure 7.

Supplementary Video S1 is related to Figure 5 (Vehicle-only group).
Supplementary Video S2 is related to Figure 5 (Olaparib-only group).
Supplementary Video S3 is related to Figure 5 (Zoledronate-only group).
Supplementary Video S4 is related to Figure 5 (Olaparib + Zoledronate group).

RNA sequencing data is accessible on website at following address (GEO Accession Number: GSE208234): https://www.ncbi.nlm.nih.gov/geo/query/acc.cgi?acc=GSE208234 (reviewer token: ypmdcowstjadfwp).

### Supplemental Figures

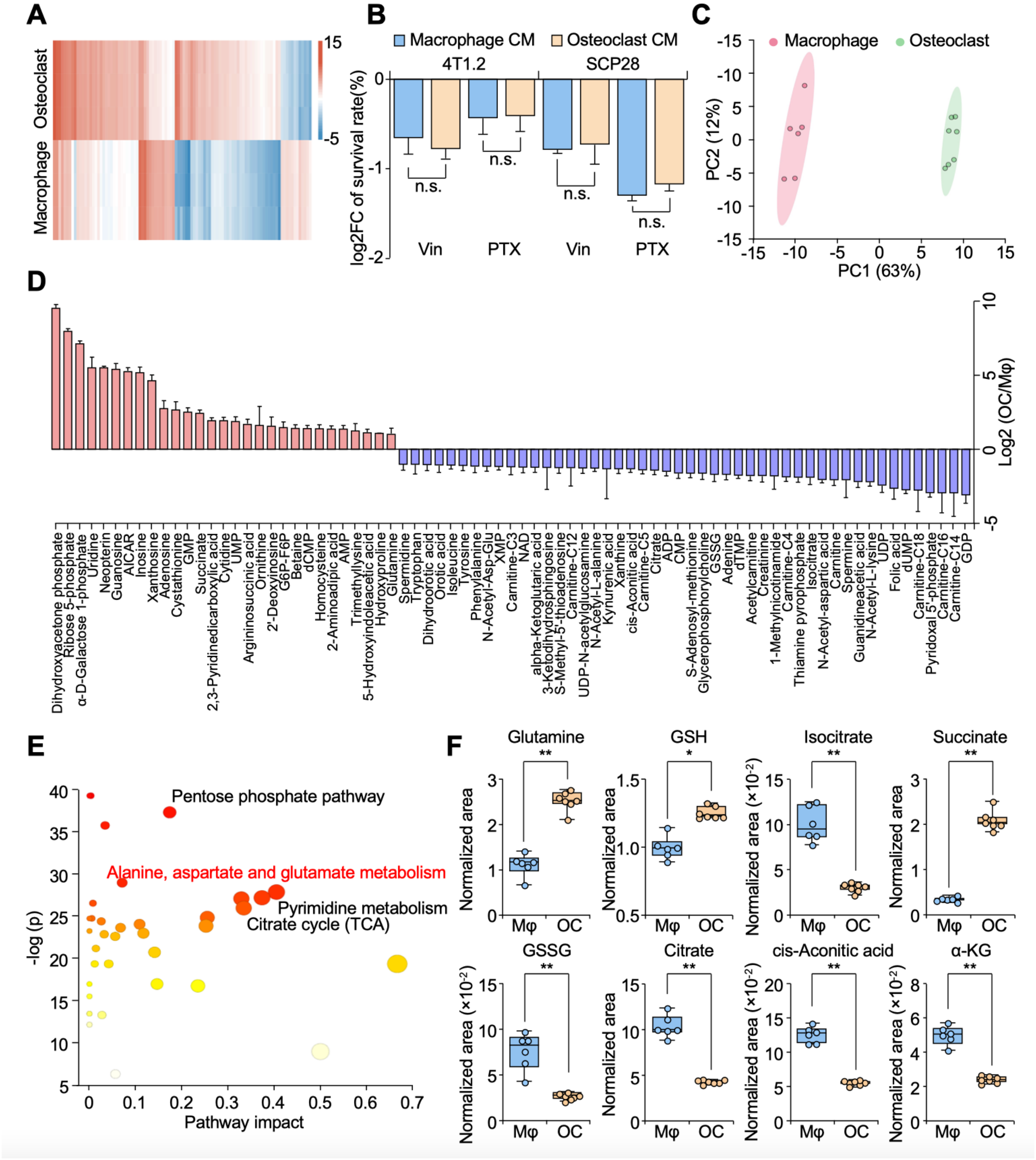


**Supplemental Figure 1, related to Figure 1. Comparison of metabolic pathways in macrophages and osteoclasts.** (**A**) Heatmap showing differentially expressed genes in macrophages and osteoclasts. (**B**) Survival rates of 4T1.2 and SCP28 cells treated with vincristine (10 nM) or paclitaxel (25 nM) in the presence of macrophage or osteoclast CM mixed with regular culture medium (ratio 1:1). (**C**) Principal component analysis of metabolomics data from macrophages and osteoclasts. (**D**) Bar graph showing significantly changed metabolites in osteoclasts compared to macrophages. Only metabolites with fold changes larger than 2 or less than 0.5 and FDR-values less than 0.05 are presented. (**E**) Metabolic pathway enrichment analysis of osteoclast versus macrophage based on significantly changed metabolites. (**F**) Normalized areas of glutamine, GSSG, GSH, citrate, isocitrate, cis-aconitic acid, succinate, and α-KG in macrophage (Mφ) and osteoclast (OC) samples. n = 6 for macrophage group; n = 7 for osteoclast group. Data represents mean ± SEM. “n.s.” means not significant, $*p < 0.05$, $**p < 0.01$ by Student’s t-test.

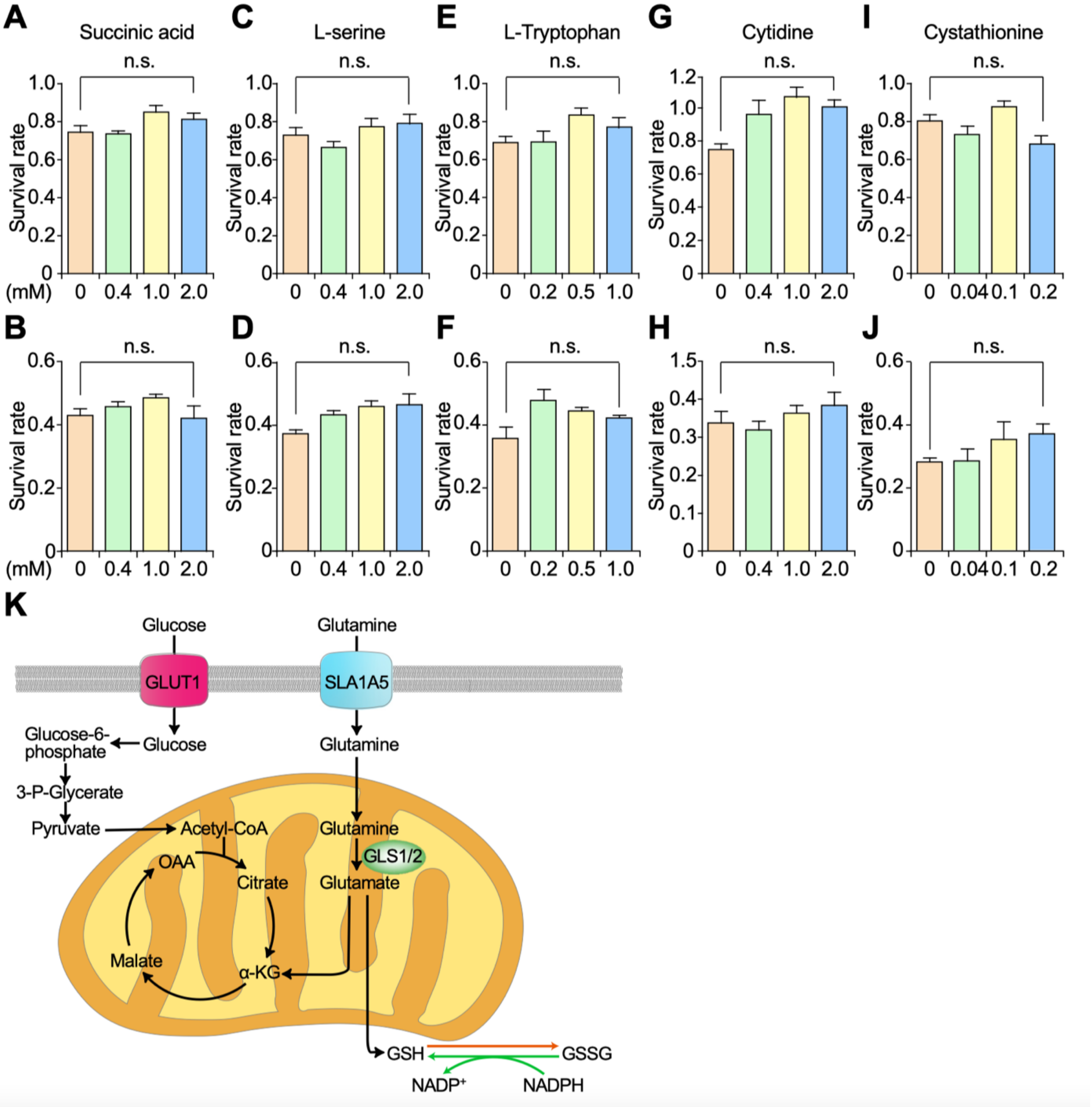


**Supplemental Figure 2, related to Figure 2. Most candidate metabolites do not induce PARPi therapy resistance in bone metastasis-prone tumor cells.** (**A-B**) The figure shows the normalized survival rates of SCP28 and 4T1.2 cells treated with either DMSO control or olaparib (25 μM) and cultured in indicated concentrations of succinic acid. SCP28 cells were used in A. 4T1.2 cells were used in B. (**C-D**) The normalized survival rates of SCP28 and 4T1.2 cells treated with either DMSO control or olaparib (25 μM) and cultured in indicated concentrations of L-serine. SCP28 cells were used in C. 4T1.2 cells were used in D. (**E-F**) The normalized survival rates of SCP28 and 4T1.2 cells treated with either DMSO control or olaparib (25 μM) and cultured in indicated concentrations of L-tryptophan. SCP28 cells were used in E. 4T1.2 cells were used in F. (**G-H**) The normalized survival rates of SCP28 and 4T1.2 cells treated with either DMSO control or olaparib (25 μM) and cultured in indicated concentrations of cytidine. SCP28 cells were used in G. 4T1.2 cells were used in H. (**I-J**) The normalized survival rates of SCP28 and 4T1.2 cells treated with either DMSO control or olaparib (25 μM) and cultured in indicated concentrations of cystathionine. SCP28 cells were used in I. 4T1.2 cells were used in J. (**K**) A simplified schematic representation of the glutamine metabolism pathway is shown. Extracellular glutamine is taken up by cells via the membrane transporter SLC1A5. Glutamine is then converted to glutamate by glutaminase 1/2 (GLS1/2). A portion of glutamate enters the TCA cycle by converting to α-KG (α-ketoglutarate), while the remaining portion is used to generate glutathione. Data represents mean ± SEM. "*n.s.*" means not significant by Student's t-test (A-J).

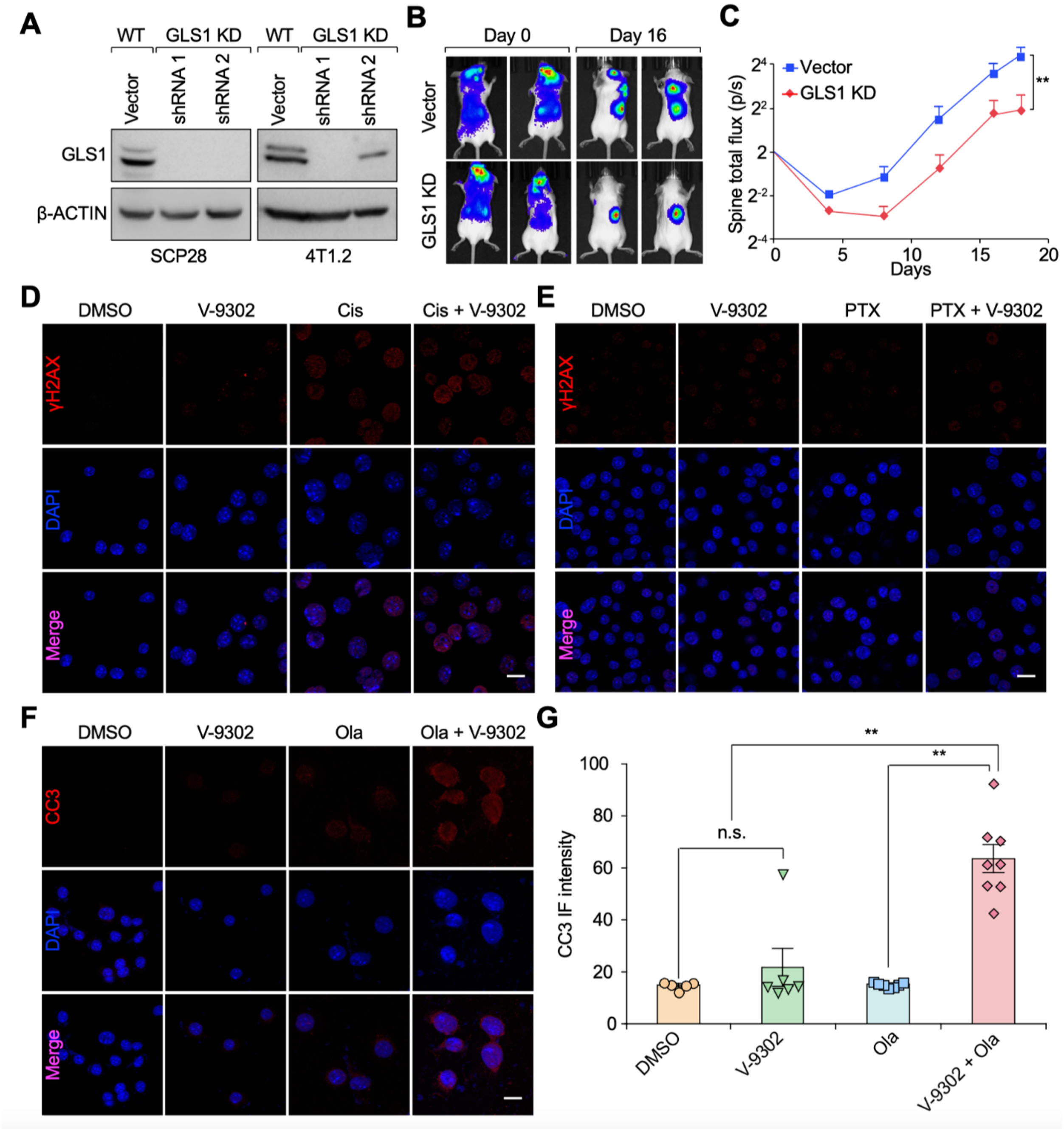


**Supplemental Figure 3, related to Figure 3. Inhibition of glutamine metabolism enhances DNA damage and cell death during PARPi treatment. (A**) Confirmation of GLS1 KD efficiency in SCP28 cells (left) and 4T1.2 cells (right) by immunoblotting. β-ACTIN was used as internal loading control. (**B**) $1{\times}10^5$ 4T1.2 cells were IC injected into 4-6 weeks old female BALB/c mice. Representative BLI images at Day 0 and Day 16 were shown. (**C**) The spine skeletal metastasis burden was quantified based on BLI imaging from the experiment in B. n =

12 per group. (**D**) 4T1.2 cells were treated with DMSO, cisplatin, V-9302, and the combination of both cisplatin and V-9302. Two days later, cells were fixed for IF staining against γH2AX (red). Nuclei were counter-stained with DAPI (blue). Scale bar, 20 μm. (**E**) 4T1.2 cells were treated with DMSO, PTX, V-9302, and the combination of PTX and V-9302. Two days later, cells were fixed for IF staining against γH2AX (red). Nuclei were counter-stained with DAPI (blue). Scale bar, 20 μm. (**F**) 4T1.2 cells were treated with DMSO, V-9302, olaparib, and the combination of V-9302 and olaparib. Two days later, cells were fixed for IF staining against Cleaved Caspase 3 (CC3) (red). Nuclei were counter-stained with DAPI (blue). Scale bar, 20 μm. (**G**) The IF staining intensity of CC3 from the experiment in F was quantified. Ola represents olaparib, Cis represents cisplatin, and PTX represents paclitaxel. Data represents mean ± SEM. "*n.s.*" means not significant. $^{**}p < 0.01$ by Student's t-test (C and G).

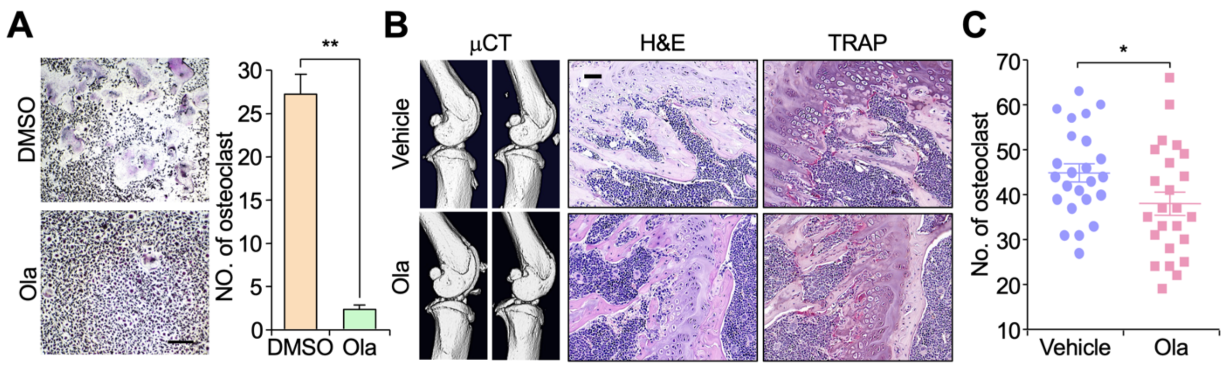


**Supplemental Figure 4, related to Figure 4. Examining potential side effects of olaparib in the bone microenvironment.** (**A**) Bone marrow cells were in vitro differentiated to generate osteoclasts using a similar protocol to that in Figure 1A. During osteoclast differentiation, the cell culture was treated with DMSO or olaparib (20 μM). Ten to fourteen days later, cells were stained with a TRAP staining kit (left panel). The number of TRAP-positive osteoclasts was quantified (right panel). Scale bar, 500 μm. n = 3 per group. (**B**) Six-week-old female nude mice were treated with either vehicle control or olaparib (50 mg/kg) for four weeks. Mice were then sacrificed for hindlimb collection and histological analysis. Representative μCT, H&E, and TRAP staining images are presented. Scale bar, 50 μm. (**C**) Quantification of TRAP-positive osteoclasts from decalcified histological bone sections of hindlimbs from mice in B. n = 24 per group. Data represents mean ± SEM. *p < 0.05 and **p < 0.01 by Student's t-test (A, C).

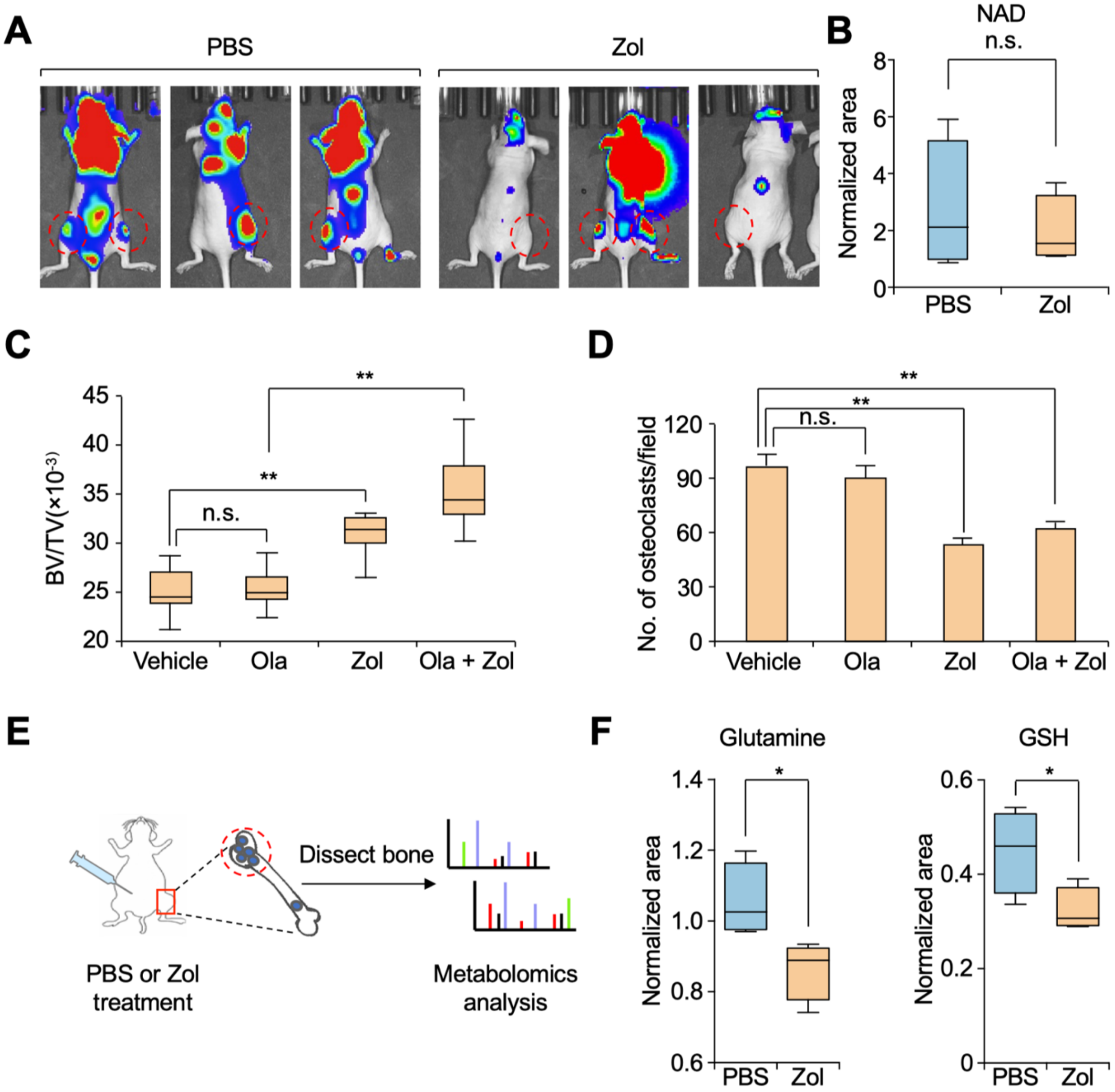


**Supplemental Figure 5, related to Figure 5. Inhibition of osteoclast activity synergistically suppressed bone metastasis progression with PARPi treatment.** (**A**) Representative BLI images of mice with bone metastases at Day 55 are shown. (**B**) Quantification of normalized area of NAD in bone metastases samples in PBS- or zoledronate-treated mice. n = 4 per group. (**C**) The quantification of bone volume (BV) *v.s.* total volume (TV) based on µCT imaging from the experiment in Figure 5I is shown. n = 12 mice per group. (**D**) The quantification of TRAP-positive osteoclasts based on TRAP staining from the experiment in Figure 5J is shown. n = 12 per group. (**E**) The workflow of analyzing metabolites in in the glutamine pathway of normal mouse bone tissues without cell inoculation. 4-6 weeks old mice were treated with either vehicle control (PBS) or zoledronate (2mg/kg) twice a week. Mice were treated continuously for about 7 weeks. The same bone areas (red-dotted area) from both groups were minced directly for sample collection and mass spectrometry analysis. (**F**) Experiment was performed as illustrated in E. Normalized areas of glutamine or GSH in PBS-treated group or in zoledronate-treated group were presented. n = 4 per group. Data represents mean ± SEM. "n.s." means not significant. $^{**}p < 0.01$ by Student's t-test (B, C, D, and F).

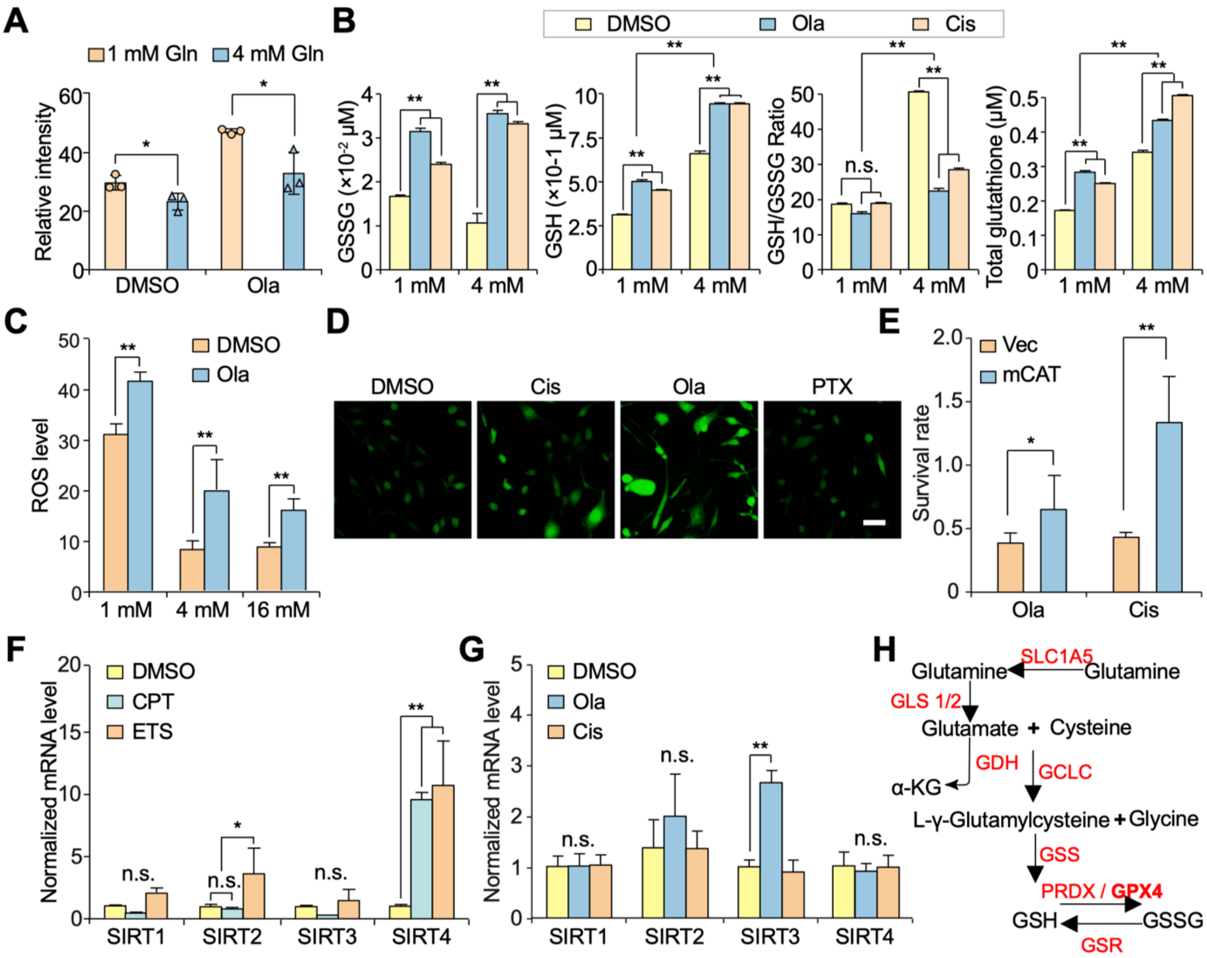


**Supplemental Figure 6, related to Figure 6. Olaparib treatment diverts glutamine to glutathione metabolism and enhances GSH to GSSG conversion.** (**A**) Quantification of ROS levels from experiment in Figure 6A. n = 3 per group. (**B**) Quantification of ROS levels in 4T1.2 cells treated with either DMSO or olaparib (25 μM) in the presence of 1 mM, 4 mM, or 16 mM glutamine. n = 3 per group. (**C**) GSH and GSSG levels, their ratio (GSH/GSSG), and the total glutathione concentration were determined in 4T1.2 cells treated with DMSO, olaparib (25 μM), or cisplatin (3 μM) in cell culture media containing either normal (4 mM) or low (1 mM) concentration of glutamine. n = 3 per group. (**D**) SCP28 cells were treated with DMSO, olaparib (25 μM), cisplatin (3 μM), or PTX (25 nM). The ROS levels in these cells were determined by a Dichlorodihyfrofluorescin diacetate (DCFH-DA) ROS assay kit. Scale bar, 50 μm. (**E**) SCP28 cells were transduced with lentiviruses to express either control Vector or mitochondria-localized catalase (mCAT). The survival rate of SCP28-Vector cells and SCP28-mCAT cells after treatment with olaparib (25 μM) or cisplatin (3 μM) for 3 days was determined by luciferase assay. (**F**) The mRNA expression levels of SIRT1, SIRT2, SIRT3, and SIRT4 in SCP28 cells treated with either vehicle control (DMSO), camptothecin (CPT) or etoposide (ETS) were determined by qPCR. *GAPDH* was used as internal control. n = 3 per group. (**G**) The mRNA expression levels of SIRT1, SIRT2, SIRT3, and SIRT4 in SCP28 cells treated with either vehicle control (DMSO), olaparib or cisplatin were determined by qPCR. *GAPDH* was used as internal control. n = 3 per group. (**H**) Schematic illustration of detailed metabolic enzymes involved in glutamine to glutathione synthesis. Data represents mean ± SEM. "n.s." means not significant. *p < 0.05, **p < 0.01 by Student's t-test (A, B, C, E, F and G).

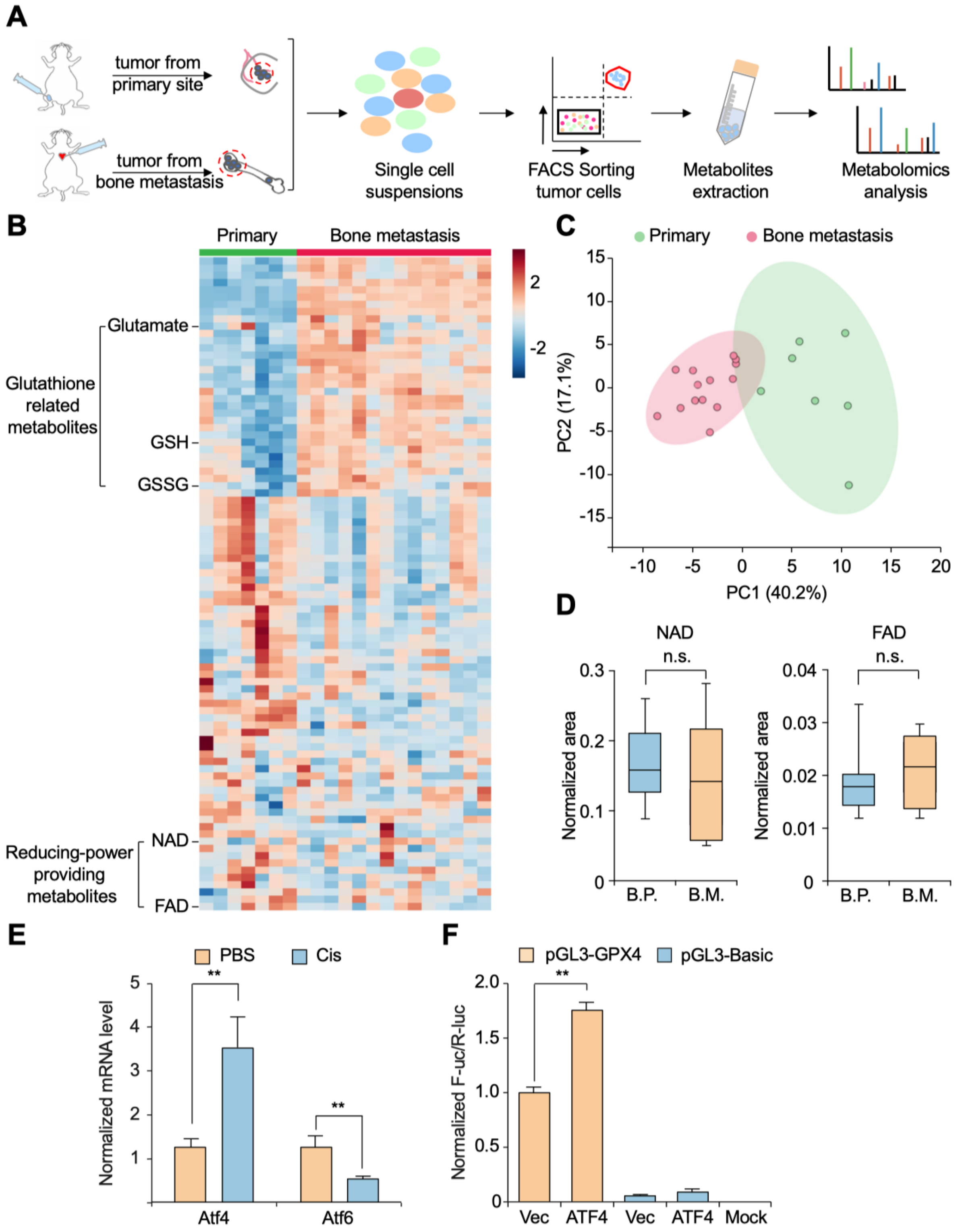


**Supplemental Figure 7, related to Figure 7. Glutathione metabolism is enhanced in bone metastasis.** (**A**) Workflow for the analysis of metabolites in tumor cells from primary or bone metastatic sites. 1×$10^5$ SCP28 cells were mammary fat pad injected or IC injected into 4-6-week-old female nude mice. Primary tumors or bone metastases which were confirmed by BLI imaging (red-dotted area) were minced for enzymatic digestion for single cell suspensions. Tumor cells were FACS sorted (by their GFP-labelling) for sample collection and mass spectrometry analysis. (**B**) Heatmap of metabolites in primary tumor cells and bone metastasis cells as determined by mass spectrometry. Primary tumor (P.T.) samples: n = 7; Bone metastasis (B.M.) samples: n = 14. (**C**) Principal component analysis of the metabolomics of tumor cells from primary sites and bone metastasis group. (**D**) Tumor cells from primary sites or bone metastases were profiled for critical metabolites. Normalized areas of NAD and FAD in cancer cells from primary tumors (P.T.) or from bone metastases (B.M.) were presented. n = 7 for primary group; n = 14 for bone metastasis group. (**E**) The mRNA expression levels of *Atf4* and *Atf6* was determined by q-PCR in 4T1.2 cells treated with PBS or 3 μM cisplatin.. *Actb* gene was utilized as internal loading control. n = 3 per group**.** (**F**) Indicated plasmids were co-transfected into 293T cells. Firefly luciferase reporter activity was measured and normalized to Renilla luciferase internal control. Data represents mean ± SEM. “n.s.” means not significant. **p < 0.01 by Student’s t-test (D, E, and F).
